# Self-supervised learning enables unbiased patient characterization from multiplexed cancer tissue microscopy images

**DOI:** 10.1101/2025.03.05.640729

**Authors:** Gantugs Atarsaikhan, Isabel Mogollon, Katja Välimäki, iCAN, Tuomas Mirtti, Teijo Pellinen, Lassi Paavolainen

**Affiliations:** Institute for Molecular Medicine Finland (FIMM), HiLIFE, University of Helsinki, Finland; iCAN Digital Precision Cancer Medicine Flagship, University of Helsinki, Finland; Department of Pathology, University of Helsinki and Helsinki University Hospital, Helsinki, Finland; Research Program in Systems Oncology, University of Helsinki, Helsinki, Finland; Finnish Cancer Institute, Helsinki, Finland

**Author notes:** Correspondence, **Corresponding authors** Correspondence to Gantugs Atarsaikhan and Lassi Paavolainen.

## Abstract

Multiplexed immunofluorescence microscopy offers detailed insights into the spatial architecture of cancer tissue. However, classical single-cell analysis approaches are limited by segmentation accuracy, reliance on predefined features, and the inability to capture spatial interrelationships among cells. We developed a hierarchical self-supervised deep learning framework that learns spatial protein marker patterns from multiplexed microscopy images by encoding the tissue at both local (cellular) and global (tissue architecture) levels. Applied to lung, prostate, and renal cancer tissue microarray cohorts, our method stratified patients into prognostically distinct groups with significantly different survival outcomes. These groupings were consistent with prior expert-driven single-cell analyses, demonstrating the validity of our approach. Furthermore, attention maps extracted from these models highlighted biologically relevant tissue regions associated with specific marker patterns. Overall, our framework effectively profiles complex multiplexed microscopy images and offers a scalable, interpretable tool for improved biomarker discovery, with potential to support more informed cancer treatment decisions.

## Introduction

Understanding the heterogeneity of cancer cell states and their surrounding tumor microenvironment (TME) is essential for patient diagnosis and treatment optimization^1,2^. Interactions between cancer cells and the TME greatly influence cancer development, progression, and response to therapy^3–5^. Previous studies have examined various components of TME, such as cancer-associated fibroblasts (CAFs)^6–9^ and tumor-associated macrophages^10–12^. Due to the TME’s complexity, studies suggest that targeting only certain parts of it is insufficient for developing optimal cancer treatments^3,13^. Therefore, a comprehensive understanding of cancer cells and their TME is needed to develop effective treatment strategies.

Multiplexed immunofluorescence (mIF) microscopy technique^14–16^ is widely used to simultaneously visualize multiple proteins in tissues. This approach is especially useful for providing detailed spatial information on cell types and protein expression levels from complex cancer tissue structures^16–22^. Typical mIF microscopy image analysis relies on cell segmentation^23,24^ and measurement of single-cell morphological and intensity features^25^, followed by various downstream analyses^16,26^. The mIF images have been employed to identify correlations between CAF types and patient survival in lung cancer^19,27^, classify cell types in head and neck squamous cell carcinoma^28^, and discover predictor cell types associated with favorable prognosis in breast cancer^29^. However, these methods rely on single-cell features, which can be limited by segmentation accuracy and sample preparation issues that yield partial or overlapping cells. Therefore, new methods are needed to fully leverage mIF images by incorporating cellular information, local and global spatial context, and cell–cell interactions.

Self-supervised learning (SSL) methods enable unbiased profiling of large image datasets without requiring expert annotations^30^. Recent advances in SSL methods for natural images have achieved impressive performance on downstream tasks^31–35^, including tumor histopathology analyses^36^. However, models pre-trained on natural images are typically not directly applicable to fluorescent microscopy images because of differences in signal distribution and the number of channels. To employ SSL methods for fluorescence microscopy images, Doron et al.^37^ and Pfaendler et al.^38^ studied DINO SSL framework^31^, and Kraus et al.^39^ proposed a channel-agnostic Masked Autoencoder (MAE) method^35^ for single cell profiling. By successfully applying SSL to cell culture image data, these studies provide a proof-of-concept that SSL can be extended to mIF microscopy images of tissue sections, which we hypothesize will yield similarly valuable insights.

In this study, we developed an SSL framework for profiling cancer tissue samples prepared as tissue microarrays (TMA) and imaged by cyclic mIF microscopy. The framework, inspired by the Hierarchical Image Pyramid Transformer (HIPT) method^36^ for whole-slide histopathology images, operates on two modular levels: local cellular neighborhoods and global tissue structures. We applied the SSL framework for characterizing cancer cohorts from lung, prostate, and renal cancers.

## Results

### Hierarchical self-supervised learning framework

We present a hierarchical SSL framework with two levels for encoding complex spatial marker patterns from mIF images (Fig. 1). This framework learns local representations of tissue that capture single-cells and their close interaction within small neighborhoods, as well as global representation that encodes overall tissue architecture, all without the need for manual annotations or segmentation. Furthermore, the framework learns multi-channel relationships in mIF images, facilitating the discovery of marker patterns that may hold significant biological relevance.

**Figure 1.**
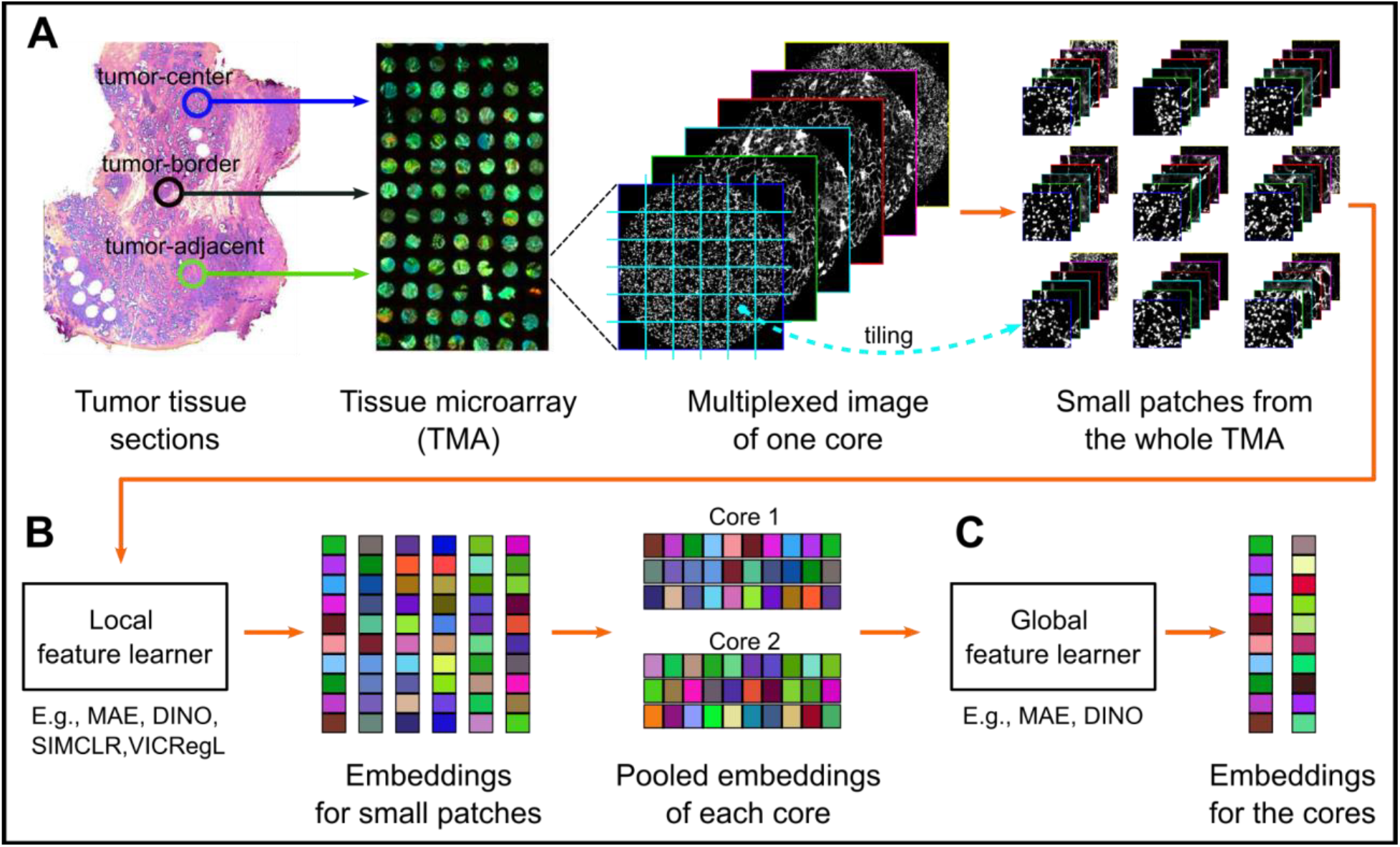
Overview of the proposed framework. **A)** Tissue microarrays (TMAs) are constructed by cutting cores from tumor tissue sections in three distinct regions: tumor-center, tumor-adjacent, and tumor-border. The TMA slides are cyclically stained with fluorescent markers and imaged to create multiplexed images. Our study cohorts consist of 7–10 TMA slides, each containing approximately 100 cores. Multiplexed images of the cores (about 4000×4000 pixels) are then divided into non-overlapping patches of 256×256 pixels, representing local cellular neighborhoods. **B)** Using the small patches, we trained a model with a self-supervised learning approach to capture local marker patterns. We tested MAE, DINO, SIMCLR, and VICRegL methods as local feature learners. **C)** To learn global feature representations, we trained a separate model using aggregated features of each core as input. We tested MAE and DINO as global feature learners.

We applied our framework to TMA images from lung, prostate, and renal cancer cohorts (Table 1). Cores with diameter of approximately 1mm were collected from three distinct regions on the tissue: *tumor-center* (includes mostly cancer cells), *tumor-border* (includes both cancer and stromal cells), and *tumor-adjacent tissue* (mostly stromal cells) (Fig. 1A). Each core image is about 4,400×4,400 pixels and was stained with six or seven markers using our in-house designed CAF panel^18,19^ (Table 1). For local feature extraction, each core image was subdivided into 256×256 pixel (80×80 µm) patches, each containing several dozen cells. Next, we applied a logarithmic transformation to the patch intensities, compressing long-tailed histograms toward normal distributions to improve SSL efficiency.

**Table 1.** Details of TMA datasets.

| Cancer Type | Lung Cancer | Prostate Cancer | Renal Cancer |
| --- | --- | --- | --- |
| **TMA core regions** | Tumor-center | Tumor-center, Tumor-adjacent | Tumor-center, Tumor-border, Tumor-adjacent |
| **Number of patients** | 319 | 299 | 237 |
| **Number of TMA cores** | 569 | 827 (585 / 242) | 1225 (501 / 481 / 243) |
| **Number of Patches** | 99,106 | 125,631 | 180,671 |
| **Number of Channels** | 6 | 7 | 6 |
| **Marker Names** | DAPI, PDGFRB, PDGFRA, SMA, FAP, PanEpi | DAPI, PDGFRB, FAP, pSTAT3, SMA, Collagen I, PanEpi | DAPI, PDGFRB, PDGFRA, SMA, FAP, PanEpi |

The local model was trained on these patch images using the Masked Autoencoder (MAE)^35^ approach with the aim of capturing hidden patterns within and between the markers. Once the local model was trained, we used it to extract feature representations of the patch images (Fig. 1B). The global model was trained with the DINO SSL method^31^, with local feature representations of cores as input feature matrices. The global model was used to extract feature representations from whole cores (Fig. 1C). Finally, patient-level feature representations were created by averaging the global feature representations across cores of the same region (tumor center, tumor border, or tumor adjacent) within each patient. Detailed description of the methodology is presented in the Methods section, and the source code of the framework is available on GitHub (https://github.com/bioimage-profiling/SSL-Multiplexed-Imaging). Evaluation of different SSL methods is presented in Supplementary methods and in (Suppl. Fig. S1).

### Encoding fluorescent marker intensity patterns via local neighborhood representations

The lung, prostate, and renal patch image datasets consist of approximately *100k, 125k*, and *180k* images, respectively. Lung and renal cancer images include six channels, while prostate cancer images contain seven channels of fluorescent markers. In the local model, we train the MAE model on each cancer dataset separately, and extract feature representation for each patch image with the aim of identifying distinctive multimarker patterns in cellular neighborhoods.

First, we checked how marker intensities are encoded in the local feature representations. We created two-dimensional UMAP embeddings from the feature representations and inspected the distribution of average marker intensities of the patches in these plots (Fig. 2A and Suppl. Fig. S4). Patches with similar average marker intensities (e.g., low PDGFRB, high Collagen I, and low PanEpi) are encoded with similar local feature representations which are mapped closely in the UMAP embedding. Specifically, on the PanEpi channel, the division between low PanEpi intensity (other than epithelial cells) and high PanEpi intensity (epithelial cells) patches is clear, although other channels do not follow such clear division in all cohorts. This indicates that the local model was able to learn patterns between multiple markers instead of only clustering based on intensities of individual markers.

**Figure 2.**
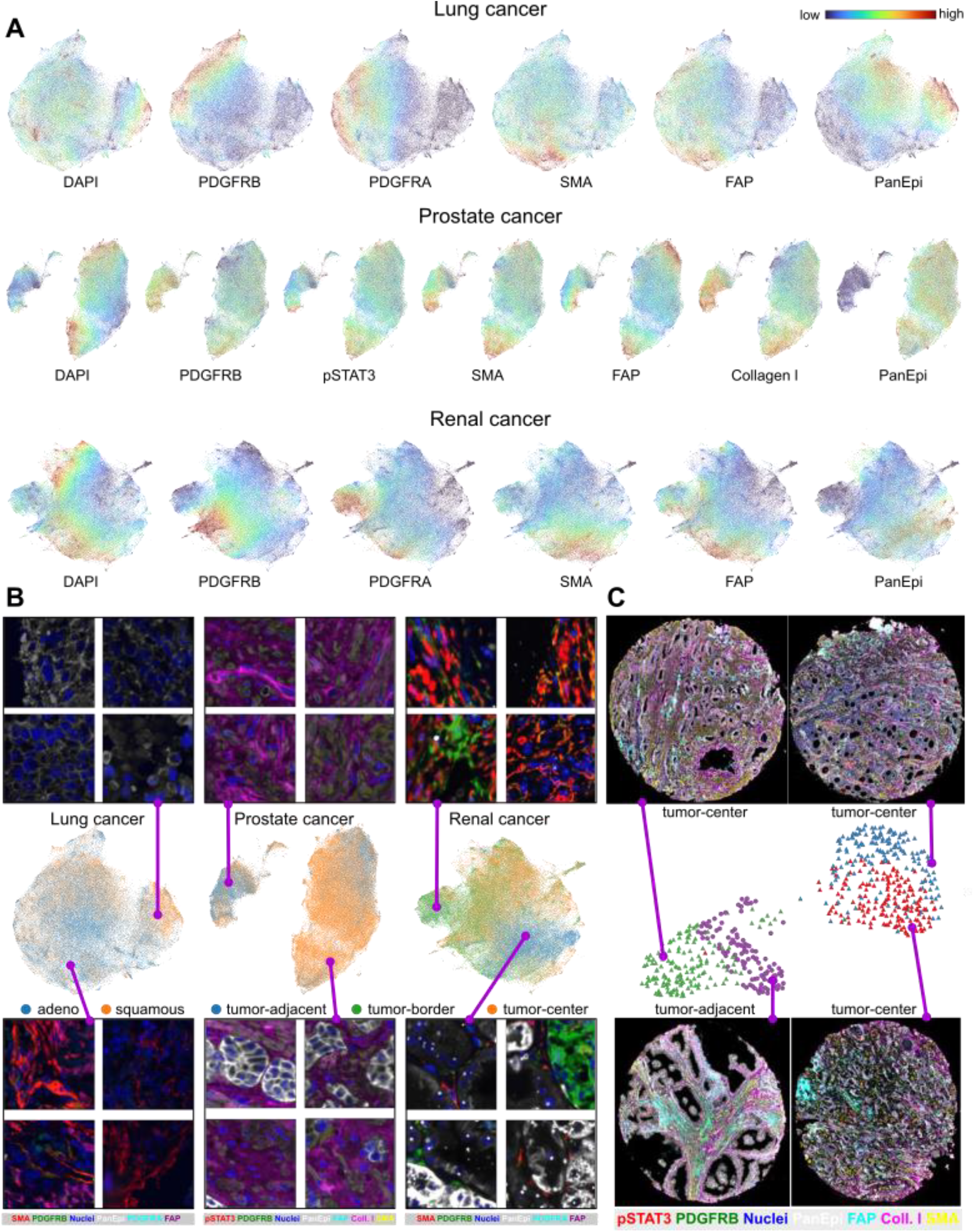
Local feature representations. **A)** UMAPs showing local (patch-image level) feature representations of three cancer cohorts by using the MAE method. The colormaps on the UMAPs show the mean intensity value of each patch image for each channel. **B)** The same UMAPs are colored by tumor region for prostate and renal cancer, and by histology for lung cancer. Selected sample images from each cohort are shown above and below the UMAPs. **C)** Feature representations of all patches from one patient in the prostate cancer cohort. Feature representations are plotted in two groups by UMAP, each containing patches from two cores. For **B** and **C** multiplexed images were mapped to composite RGB images for visualization.

Next, using the same UMAP embeddings, we confirmed that patch feature representations are able to distinguish between core regions on the prostate and renal cancer datasets (Fig. 2B). In addition, patches from different regions share considerable overlap in the embedding space with low silhouette score (0.09 for the prostate cohort and 0.01 for the renal cohort). We hypothesize that the feature representations are primarily distinguished by structural markers: nuclear and epithelial markers. Then, these features are subdivided by combinations of CAF-specific markers: PDGFRB, PDGFRA, SMA, and FAP. All cores in the lung cancer cohort were sampled from the tumor tissue, thus we studied the patch embeddings by sample histological subtypes that are visibly divided in the UMAP.

Lastly, we checked all local feature representations from one randomly chosen patient with prostate cancer (Fig. 2C). The cohort includes one tumor-adjacent core and three tumor-center cores from this patient. The UMAP embedding shows that local feature representations from the same core are more similar than from other cores. Visual inspection showed that the cores with higher epithelial and pSTAT3 markers are plotted in one group, and the other group has higher nuclear and stromal signals. Overall, local models successfully encoded multi-channel patch images into patch feature representations showing distinct characteristics.

### Global feature representations of cores distinguish between sample types

For each core, we pooled the local feature representations and provided them as a single input to the global model. Thus, the dataset sizes for each cancer cohort are around 500, 800, and 1200 for the lung, prostate, and renal cancer cohorts, respectively. We trained the global models with the DINO SSL method and extracted features from each core (Fig. 3A). In the case of lung cancer dataset, visually similar cores are grouped together in the UMAP space (Fig. 3B). The model also encoded the majority of the adenocarcinoma samples with similar feature representations. Prostate and renal cancer feature representations were separated by the core regions. We measured the overall separation of the feature representations using the normalized Mean Pairwise Distance (nMPD), which ranges from 0 to 1: 0 indicates that all embeddings are identical, while 1 indicates that each embedding forms its own distinct cluster.

**Figure 3.**
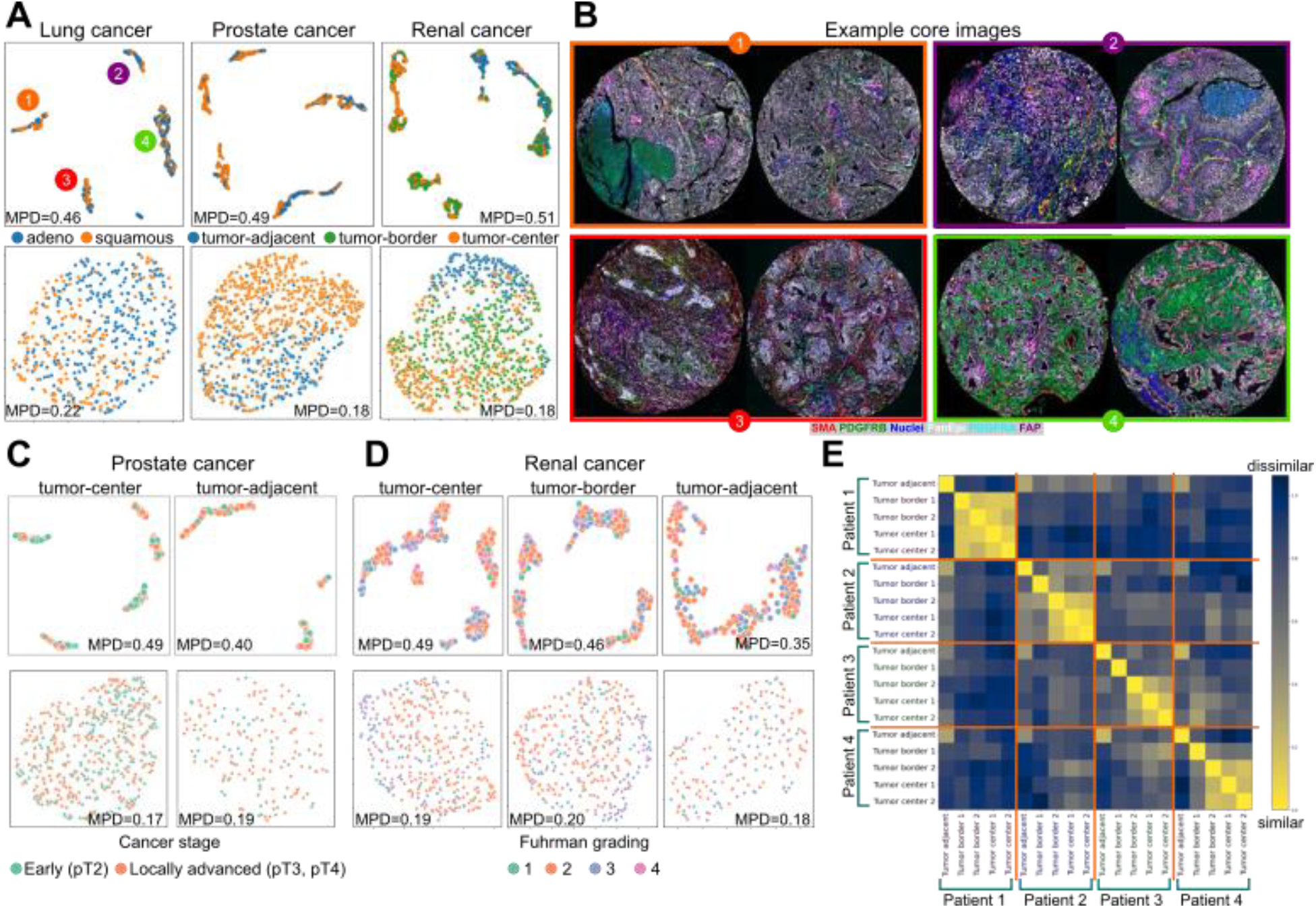
Global feature representations. For each representation, the normalized Mean Pairwise Distance (nMPD) score is calculated. The nMPD score is close to 0 when the embeddings are almost identical, and close to 1 when the embeddings are highly diverse and lack any formed clusters. **A)** Top row: global feature representations (core-level) for each cancer type. Bottom row: Features averaged per-core from all local feature representations. UMAP embeddings are colored by histology for lung cancer, and by core region for prostate and renal cancer. **B)** Example cores from different clusters defined by the global features in the lung cancer cohort. The multiplexed images were mapped to composite RGB images for visualization. **C, D)** Top row: region-wise global feature representations from the model. Bottom row: region-wise per-core averaged local feature representations. Dots in the UMAPs are colored by cancer staging for prostate cancer (C), and Fuhrman grading for renal cancer (D). **E)** Cosine distances between global feature representations from four randomly selected renal cancer patients. Low cosine distance means the features are similar to each other.

As a baseline result to compare with the global model outcome, we created core embeddings by averaging the local feature representations (bottom row of Fig. 3A). There were negligible distinctions between cores using this approach with nMPD scores from 0.18 to 0.22 depending on the cohort. In contrast, our proposed approach was able to learn and enhance minor distinctions in the local feature representations with nMPD scores from 0.46 to 0.51. To study each region independently, we created UMAP embeddings from the global feature representations of the regions separately (top row of Fig. 3C,D). In general, tumor-adjacent feature representations are less distinctive (nMPD=0.40 for prostate, nMPD=0.35 for renal) compared to those from other regions (nMPD=0.48–0.49, depending on the cohort and region), yet still more distinct than the averaged local embeddings (nMPD<=0.20). When annotated with patients’ clinical information, none of these groupings showed visually clear clusters suggesting that the extracted features capture more nuanced or deeper patterns than related to individual metadata fields (Suppl. Figs. S5-S7).

Finally, we calculated cosine distance between global feature representations of randomly selected patients with renal cancer to study similarity of the feature representations (Fig. 3E). Mean cosine distance (±standard deviation) between all pairs was 0.80±0.17. As expected, feature representations of tumor-center cores from the same patient were more similar to each other (0.17±0.13) than to those from different patients (0.82±0.05). Tumor-adjacent cores were uniquely dissimilar to tumor-center (0.85±0.14) and tumor-border (0.82±0.15) cores, even within the same patient, and were only loosely correlated with tumor-adjacent cores from other patients (0.61±0.23). The similarity of tumor-border cores varied: in some patients, tumor-border cores were correlated with tumor-center cores (Patient 1 and 2 in Fig. 3E), while in others, they showed no correlation with any region (tumor border 1 in Patient 3 in Fig. 3E).

### Feature representations uncover patients with clearly distinct prognosis

Finally, we created patient-level feature representations by averaging the global features for each patient, separately for each region: tumor-adjacent, tumor-border, and tumor-center due to the heterogeneity between regions. Then, the k-means clustering method was used for patient categorization. The number of clusters was determined by the elbow method^40^, which coincidentally gave *k=3* for all experiments. We associated the patient groups with overall survival for lung cancer, recurrence-free survival after radical prostatectomy for prostate cancer, and cancer-free survival for renal cancer. Survival probabilities were confirmed using Kaplan-Meier plots and validated with hazard rates from the Cox proportional hazards model (Fig. 4, Suppl. Fig. S8). Detailed clinical information on all patient clusters can be found in the Supplementary materials (Suppl. Table. S2-S7).

**Figure 4.**
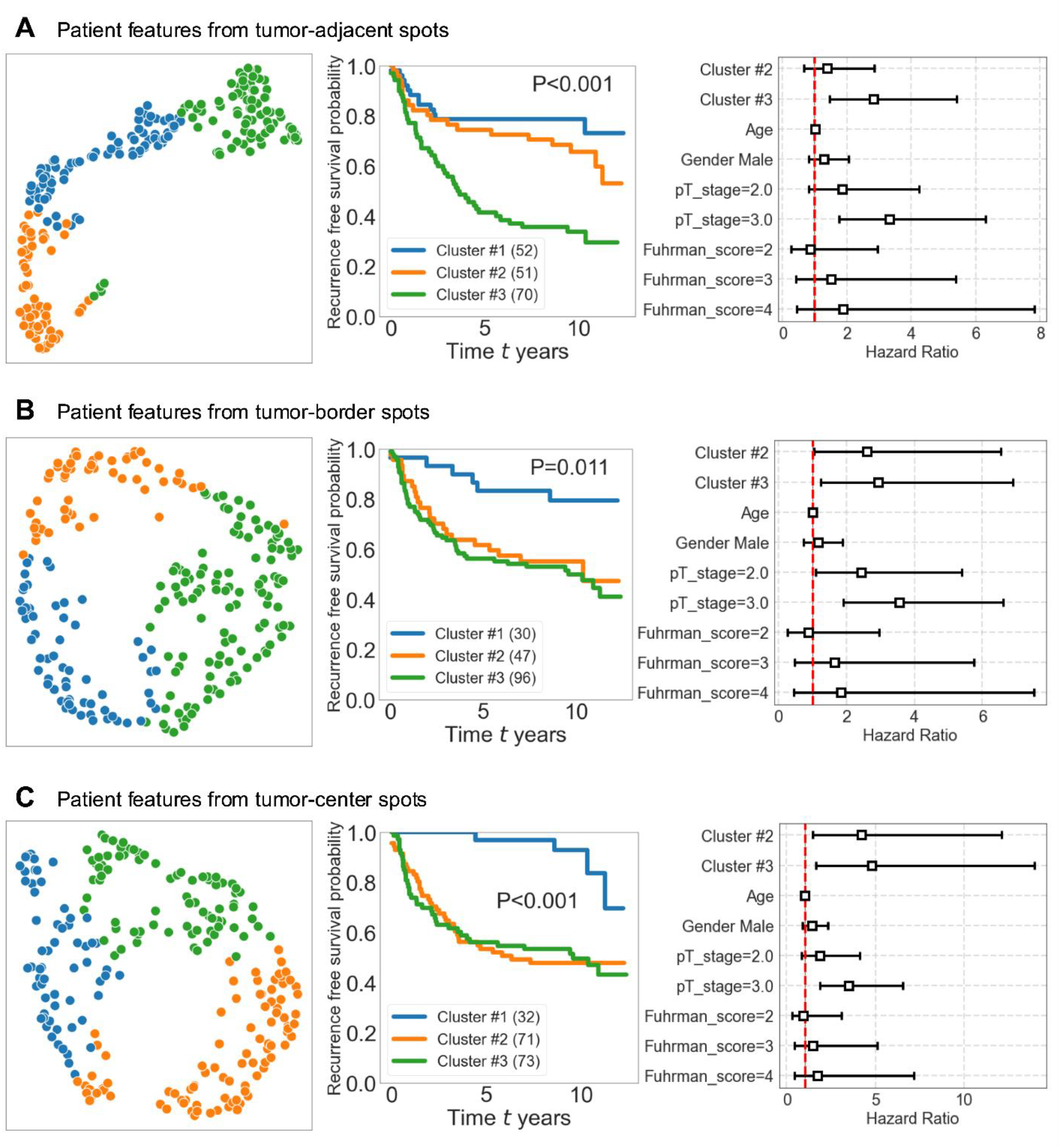
Cancer-free survival analysis of clear cell renal cell carcinoma stratified by patient-level feature representations. Patient-level feature representations were created individually by averaging all global level features from the same region: **A)** Tumor-adjacent **B)** Tumor-border **C)** Tumor-center. We clustered the original high dimensional feature representations with k-means clustering using the elbow method to determine the number of clusters. P-values were calculated using the Multivariate Log-Rank Test. The color code in UMAP graphs presents the cluster ID. We studied patient survival or recurrence using Kaplan-Meier plot and calculated the hazard rate using Cox proportional hazard regression.

We performed survival analyses for patient groups from clear cell renal cell carcinoma (ccRCC) (Fig. 4). Using tumor-adjacent cores of ccRCC, three distinct groups were identified by clustering the feature representations. Patients in the cluster#3 had the lowest survival probability. In this cluster, nearly 60% of the patients experienced relapse after initial treatment (19% in cluster#1 and 32% in cluster#2), of which 51% occurred in the lung (Suppl. Table S4). On the other hand, most patients in cluster#1 had a good prognosis with the smaller tumor and without relapse. In similar manner, patients in tumor-border cluster#1 and tumor-center cluster#1 had a good prognosis. There was no difference between genders within the clusters, except in the tumor-center cluster#1, 69% were men (52% in cluster#2 and 45% in cluster#3). Even though there was no significant difference in cancer aggressiveness indicated by the Fuhrman score, the model identified clusters with distinct prognoses demonstrating the capability of the proposed framework to extract information from mIF microscopy images not encoded in the Fuhrman score.

The lung cancer cohort was separately studied between the two main subtypes: adenocarcinoma and squamous cell carcinoma. Our framework identified three different clusters. Cluster#3 from lung adenocarcinoma patients showed significantly lower survival probability (Suppl. Fig. S8A,B). This cluster consists of more *current smokers* and have poorer *performance status scores* according to the World Health Organization (PS-WHO)^41^ criteria compared to the other two clusters (Suppl. Table S2). In the lung squamous cell carcinoma patient subgroup, cluster#3 had the lowest and cluster#1 had the highest survival probability. Patients in these groups have similar clinical background, except cluster#3 patients tend to be younger (67 or lower), and more than half have the PS-WHO score of 1 (fully ambulatory with some symptoms). In comparison, more patients did not have any symptoms (PS-WHO=0) in cluster#1 and cluster#2. At the last recorded time of follow-up, 87% of the cluster#3 patients were deceased compared to 60% of patients in the cluster#1.

For prostate cancer patients survival analysis, we considered time from radical prostatectomy (surgical removal of prostate) until status at the time of follow-up or year of cancer-related death. The framework again identified three distinct clusters from the feature representations of both tumor-center and tumor-adjacent regions. Survival probability using tumor-adjacent samples show a large difference between cluster#1 and cluster#3 patients (Suppl. Fig. S8C). More patients in tumor-adjacent cluster#3 have higher biopsy Gleason and RP/TURP Gleason scores, which may be associated with poor prognosis. The results from tumor-center samples show that patients from cluster#1 tends to have better survival probability than patients in other clusters, though the difference is not statistically significant (p=0.087, Suppl. Fig. S8D).

### Interpretation of identified patient clusters by CAF associations and attention maps

Lastly, we studied the patient clusters discovered by the hierarchical SSL method by comparing those to previously published cancer-associated fibroblast (CAF) subsets (Suppl. Table S1), defined using single-cell segmentation and expert-annotated cellular marker positivity^18,19^. CAF enrichments were defined as the ratio of CAF type cell count and the total CAF cell count. In the lung cancer cohort, we found that CAF7 (PDGFRB+, PDGFRA-, SMA+, FAP+) is enriched in patient groups with poor prognosis, and CAF13 (PDGFRB+, PDGFRA+, SMA+, FAP-) is enriched in patient groups with good prognosis (Fig. 5A, Suppl. Fig. S11). These findings match perfectly with previously published single-cell results^19^. In the renal cancer cohort, CAF associations vary between core regions, although CAF13 is associated with good prognosis regardless of the region (Suppl. Fig. S9). CAF4 (PDGFRB+, PDGFRA-, SMA-, FAP-) is enriched in the patient group with poor prognosis in the tumor-adjacent region. CAF8 (PDGFRB-, PDGFRA+, SMA-, FAP-), CAF12 (PDGFRB+, PDGFRA+, SMA-, FAP-) and CAF13 (PDGFRB+, PDGFRA+, SMA+, FAP-) are all associated with good prognosis in tumor-border and tumor-center regions whereas CAF2 (PDGFRB-, PDGFRA-, SMA-, FAP+) and CAF3 (PDGFRB0, PDGFRA-, SMA+, FAP+) are associated with a group of patients with poor prognosis. CAF6 (PDGFRB+, PDGFRA-, SMA-, FAP+), and CAF7 (PDGFRB+, PDGFRA-, SMA+, FAP+) are enriched in the tumor-center region, consistent with previous single-cell analysis^18^.

**Figure 5.**
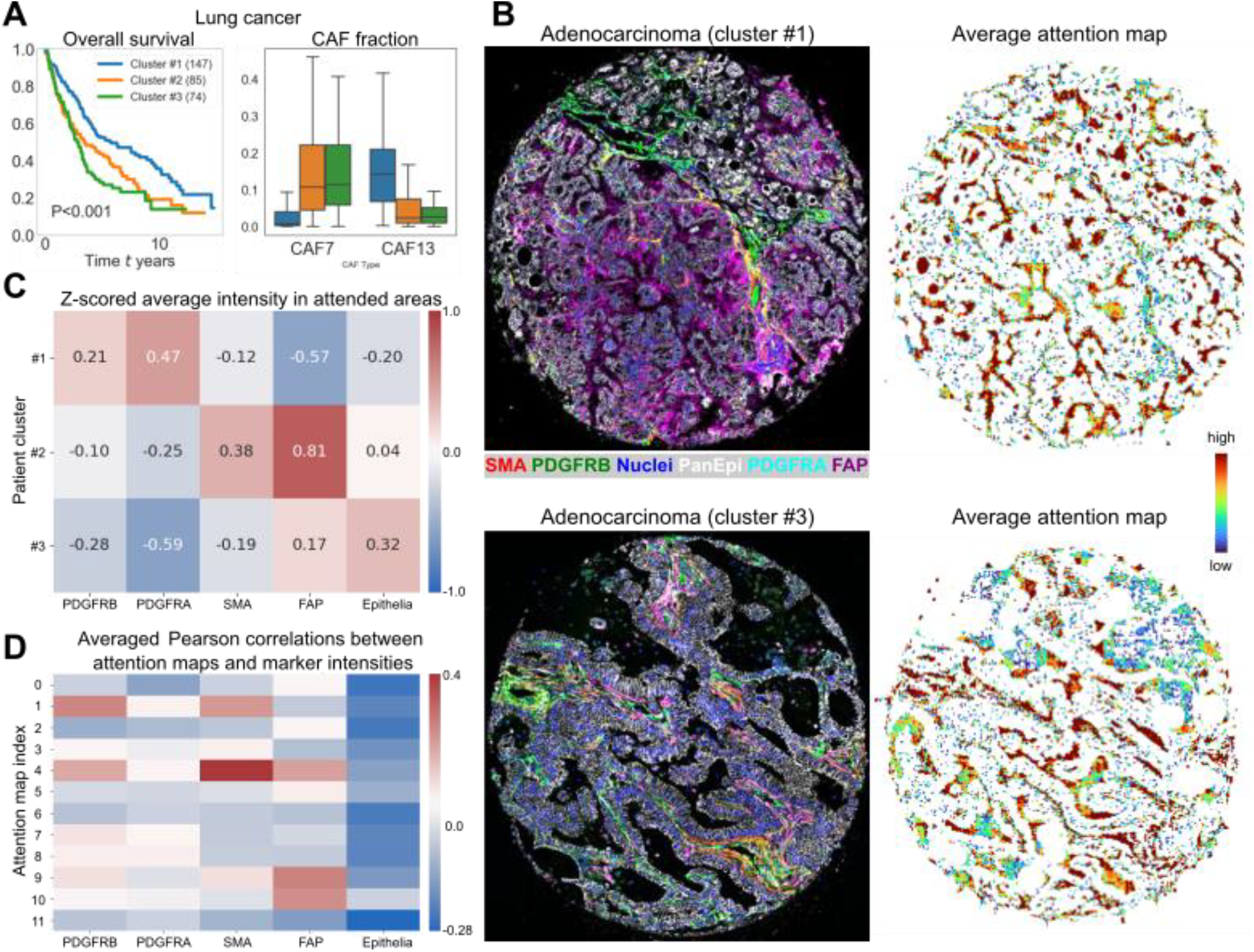
Associations between patient clusters and cancer-associated fibroblasts (CAFs) in the lung cancer cohort. **A)** Kaplan-Meier plot for patient clusters determined from patient-level feature representations, with the corresponding CAF ratio (ratio of CAF subtype and all CAFs) in each cluster for the lung cancer cohort. Subsets CAF7 (PDGFRB+, PDGFRA-, SMA+, FAP+) and CAF13 (PDGFRB+, PDGFRA+, SMA+, FAP-) were shown to be prognostic in an earlier study. **B)** The original composite images of cores, their corresponding average attention maps from good prognosis (cluster #1) and poor prognosis (cluster#3) groups. Attention maps show regions that are concluded important by the model. **C)** Normalized average intensity of the most attended areas, as defined by values above 75th percentile per attention map, across all cores in each cluster for every fluorescent channel. **D)** Pearson correlation between the attention maps and each fluorescent channel in the most attended regions, averaged across the dataset.

An important advantage of using a ViT architecture is the ability to visualize attention maps, which highlight the regions in the input images where the model focuses on. A ViT model can generate multiple attention maps depending on the number of self-attention heads employed. We used the ViT-Base architecture with 12 attention heads. Average attention maps extracted from selected lung cancer images show that the model does not focus on the abundant epithelial or nuclear cells, but rather highlights other markers (Fig. 5B, Suppl. Fig. 10). Individually, each attention head highlights a different combination of fluorescent markers (Fig. 5D). For instance, attention map #1 correlates with CAF5, whereas #4 correlates with CAF7. Attention maps #9 and #10 correlates most strongly with the FAP marker. Moreover, none of the attention maps correlate with the abundant PanEpi marker.

To identify what the model highlights within each patient group in the lung cancer cohort, we analyzed marker values inside the “most attended areas” defined as the pixels with values above the upper quartile of each individual attention map. We calculated the average intensity of each fluorescent marker within these attended areas (Fig. 5C). The good prognosis patient group (cluster#1) had higher intensities of PDGFRB and PDGFRA channels compared to the poor prognosis groups. On the other hand, the poor prognosis clusters (#2 and #3) showed higher intensities of SMA, FAP, and PanEpi markers within the attended areas.

## Discussion

We developed a hierarchical SSL framework to uncover unbiased representations from mIF images of cancer tissue sections. This framework includes local and global SSL models that extract both local and global representations from mIF tissue images. The global features are further aggregated into patient-level representations to study associations with clinical data. Our approach enables exploratory protein association analysis in tissue mIF microscopy images without any human bias or annotations.

We applied our framework to lung, prostate, and renal cancer cohorts. We showed that local features generally encode patterns of fluorescent marker intensities. This is beneficial, as the model can recognize multi-marker patterns. Our cohorts include markers for nuclei and epithelia (additionally stroma in prostate cancer cohort), which show the general tissue structure (Fig. 2A). The UMAPs of local features show that patches with high amounts of nuclei and epithelial cell signal are projected together, whereas patches with high stromal intensity were projected separately from the high epithelial marker intensity enabling distinction between epithelial and stromal cells without segmentation. Other markers exhibited more variability in patches, which provided various local marker patterns to explore on the global level.

Even though trained without any supervision, the models revealed clusters of patients that present significantly different prognosis in the survival analysis (Fig. 4 and Suppl. Fig. S8). For instance, cluster#1 discovered from the renal cancer cohort tumor-center cores showed significantly better prognosis compared to the other two discovered clusters (Suppl. Fig. S9). Inversely, cluster#3 from tumor-adjacent cores in the same cohort showed significantly worse prognosis. In the prostate cancer cohort, identified cluster#1 and cluster#3 from both tumor-adjacent and tumor-center cores indicated different survival probabilities, although the difference was not statistically significant.

Our findings from the lung cancer cohort were in-line with previous work by Pellinen et al.^19^. They showed, using classical single-cell analysis approaches, that CAF7 and CAF13 subcategories were associated with poor and good prognosis, respectively. The clusters that are automatically discovered with the hierarchical SSL framework showed that the good prognosis patient group has low CAF7 and high CAF13 fractions whereas the poor prognosis patient groups have opposite CAF7 and CAF13 fractions. In the renal cancer cohort, we found that in general good prognosis groups are associated with CAFs with FAP-, and poor prognosis groups are associated with CAFs with FAP+. This observation validates previous work on clear cell renal cell carcinoma study^18^.

Moreover, the model focused on those specific CAFs through attention maps that highlight relevant areas within the core images, indicating areas where the model is focusing on. The ability to visualize attention maps from the ViT models greatly enhances the interpretability of our framework. In the lung cancer cohort example, good prognosis groups generally have higher amounts of PDGFRB and PDGFRA markers, which are usually associated with better outcomes. On the other hand, bad prognosis groups have significantly higher levels of the FAP marker, which is typically linked to poor prognosis^18^. Moreover, individual attention heads correlated with various CAF combinations and fluorescent markers. In particular, poor prognosis–related CAF7 or FAP-positive combinations were the most prominent.

One limitation of our framework is that the models are separately trained on specific cancer cohorts and may not generalize well to unseen data. To analyze different datasets, the models should be trained from scratch with the new dataset. However, as the mIF images are extremely complex with practically an unlimited number of different fluorescent marker combinations, currently no fully generalized foundation model exists for mIF image data. Furthermore, our datasets include fluorescent markers specifically adapted for studying CAFs within the tumor microenvironment, therefore, the biological interpretations are limited to CAF cells. Potential directions for improvement include developing a channel-adaptable framework, that is capable of handling images with an arbitrary number of channels, for foundational pretraining, and fine-tuning these models with specific clinical information, such as disease severity, to assess how the model and attention maps evolve in response to this additional supervision.

In conclusion, this study demonstrates the potential of self-supervised learning for the analyses of mIF images of cancer tissue cohorts. We have developed a framework that effectively captures complex fluorescent marker patterns in tumor samples without supervision, and enables associating these patterns with clinical information, such as survival probability.

## Methods

### Samples

Our multiplexed image dataset is prepared in-house using human cancer tissue samples. We used the following renal (kidney), lung, and prostate cancer cohorts in this study.

#### Renal cell carcinoma cohort

Formalin-fixed, paraffin-embedded (FFPE) surgical specimens from 236 treatment-naive patients with localized (N0M0) ccRCC who underwent partial or radical nephrectomy at Helsinki University Hospital between 2006 and 2013 were utilized in this study. Patients with distant metastases (M1), regional lymph node metastases (N1), a history of kidney cancer, or multiple kidney tumors at diagnosis were excluded. These specimens and the corresponding tissue microarrays (TMAs) had been previously collected and constructed by Pellinen et al.^18^. The TMAs included two cores from the central tumor (1.0 mm), two from the tumor border, and two from adjacent benign kidney tissue. Digital slides were scanned and annotated for TMA construction. Two pathologists selected representative tissue blocks. Clinical and pathological data—including TNM stage, tumor grade (4-tier Fuhrman), sarcomatoid/rhabdoid differentiation, necrosis, and survival outcomes—were collected from medical records. The clinical association analyses were carried out including only the clear cell carcinoma histology (n=179). The follow-up cutoff date was September 9, 2019.

#### Lung carcinoma cohort

The FFPE TMA cohort BOMI1 construction^42,43^ and the mIF staining/imaging^19^ has been previously described. Briefly, the cohort consisted of non-small cell lung carcinoma patients surgically resected at the Uppsala University between 1995 and 2005. All specimens were reviewed by board-certified pathologists and representative areas were encircled on tissue slides before cores were taken from corresponding tissue blocks and incorporated into recipient blocks. All tumor samples were included in duplicates (2 × 1 mm tissue cores). Analysis was performed on 319 patient cases with high quality mIF image data produced by Pellinen et al.^19^. Tumor histologies represented adenocarcinoma (n=168), squamous cell carcinoma (n=111), and other (n=40).

#### Prostate cancer

The prostate cancer cohort consisted of 274 patients with Grade group 2-4 localized cancer treated by open or robot-assisted-radical prostatectomy at Helsinki University Hospital (HUS) between the years 1992 and 2015. Tumor center and adjacent benign samples were from 274 and 220 patients, respectively. Clinical association analyses were carried out with 173 and 148 patients for tumor center and adjacent benign areas, respectively. This cohort’s TMA design has been previously characterized by Lehto et al.^44^.

### Cyclic multiplexed immunofluorescence and imaging

The widefield microscopy imaging was done with two rounds of staining of fluorescent markers, and washing the cores for the next staining round. In total, we used six fluorescent markers for renal and lung cancer TMAs, and seven fluorescent markers for prostate cancer TMAs. In addition, the nuclei marker is used in both rounds of imaging to enable image registration between cycles.

### Dataset

Table 1 summarizes our datasets. In total, we have approximately 2,500 cores from around 850 patients across three cancer cohorts. In the renal cancer cohort, in total 3 to 6 cores from tumor-adjacent, tumor-border, and tumor-center areas from each patient were cropped. In total 3 or 4 cores from tumor-adjacent and tumor-center regions were collected from each patient in the prostate cancer cohort. In the lung cancer cohort, each patient has 1 to 2 cores from the tumor-center area.

Core image dimensions (*∼4500×4500 px*) do not fit into GPU memory during the training process when using a large batch size required by self-supervised learning methods. Thus, we cropped the TMA images into small patches of *256×256 px*, while preserving the number of channels. Small patches were treated as local cellular neighborhoods of a few dozen cells. Then, we converted the patches to logarithmic scale to manage the wide range of raw pixel values. In all our core images, the raw values are in the range of 0–65535 (16-bit). By converting to logarithmic scale, we avoided specifying any cut-off thresholds and were able to transform the long-tail histogram. We hypothesized that any unnecessary artifacts would have similar feature representations from the self-supervised learning phase, so it would be fairly simple to discard artifacts in post-processing, thus image artifacts were not removed beforehand. The details of the datasets are presented in Table 1.

### Local feature learner

Fig. 1 summarizes our proposed framework. Our framework consists of two levels of self-supervised training and aggregation to patient level features. As a local feature learner, we evaluated four methods: SimCLR, DINO, MAE, and VICRegL. SimCLR has two encoders to process two different transformations of the same input, each producing a feature representation. It then calculates contrastive loss that minimizes the distance with a weighted cross-entropy loss between feature representations, and updates the encoder model weights. DINO has teacher and student encoders, and the student encoder tries to learn to match the output of the teacher encoder, and updates the weights accordingly. The teacher encoder is updated with regard to the student encoder. VICRegL has three main objectives: increasing variance between images, maintaining invariance between feature representations of the same image, and minimizing covariance between feature dimensions to avoid redundant features. In MAE, a large portion of the input image is masked (70-90%), and only visible parts are fed into an encoder. Then the encoder output is used to reconstruct the image with a decoder. The model weights are updated with a reconstruction loss between the original image and the reconstructed image.

We used ViT^45^ models as encoders for DINO and MAE approaches. Specifically, we used the ViT-Base model architecture with 12 attention heads. Patch embedding module was modified to accept six channel images in case of lung and renal cancer datasets, and seven channel images in case of prostate cancer dataset. MAE uses a similar VIT decoder network that outputs the reconstructed image. In SimCLR and VICRegL approaches we used ResNet-50^46^ architecture. ResNet-50 has 50 convolutional layers with residual connections between layer blocks. The first convolutional layer was modified for input images with 6 or 7 channels. The size of feature representations from ViT-Base models is 768, compared to 2048 from ResNet-50 models. In all our frameworks, our code is based on implementations of LightlySSL^47^.

The original augmentations for the self-supervised approaches are designed for natural images, and are not optimized to fluorescence microscopy images. We removed all of the color jitter transformations (*random brightness, contrast, saturation, and hue*), and included horizontal flip, vertical flip, and rotation. Cropping and gaussian blurring were kept. All the augmentations were randomly applied with 0.5 probability. We kept the augmentations of original images as strict as possible to prevent unrealistic transformations in fluorescence microscopy images and to learn more from the existing data. After training the local model, we fed the patches once more with only center cropping augmentation and extracted the feature representations. Feature representations are class tokens from ViT models or flattened and pooled last layer features from ResNet-50 models. Lastly, feature representations from the same dataset and the same method were L2 scaled.

All approaches were trained using one node of the LUMI supercomputer (https://lumi-supercomputer.eu/) that has four AMD MI250x GPUs. The used batch size was 512 for all methods and datasets. For MAE, the masking parameter was set to 75% of the input. Model training time varied by the approach and dataset, although it took around 12 hours to train 500 epochs with the renal cancer dataset. However, we stopped the training once UMAP visualization of feature representations started showing signs of overfitting, that is patch images from each core started to group together.

### Global feature learner

The global model takes the local feature representations of patches cropped from the same core image as inputs. We perform the following aggregation to create a single input for each core. Suppose a core has *N* patches, and the local feature representations are *F*_*k*_ ∈ ℝ^*D*^, where *k* = 1,2, …, *N*, where *D* is the length of the feature vector. We concatenate these feature representations into a single matrix *A* = [*F*_1_, *F*_2_, …, *F*_*N*_] ∈ ℝ^*NxD*^. The height *H* and width *W* dimensions are created from the *N*, as *A* ∈ ℝ^*NxD*^ → *A* ∈ ℝ^*HxWxD*^to serve as input for the global model. We chose *N* = 256 = 16*x*16 to create aggregated feature-images of 16*x*16*xD*, applying zero padding if the patch count did not reach 256. This step ensured that the global model can learn relationships among the local feature representations to derive global features for the entire core. This is inspired by a similar technique proposed by Chen et al.^36^ for H&E stained histopathological images. Using the aggregated images, we trained the ViT-Base network with DINO and MAE approaches separately. We used 1×1 convolutions during patch embedding, effectively treating each pixel of the aggregated images as a separate token. The augmentations of the DINO approach were similar to that of the local model, except global random crop size was set to 14 and local was 8. For MAE, the mask ratio was the same 0.75 as in the local model. Both frameworks were trained for 100 epochs that took around an hour. After training, features representing each core were extracted and L2 normalized.

### Patient-level features

Cores from different regions are drastically different from each other even if the cores are from the same patient. Thus, we aggregated core feature representations to patient level by averaging each region separately. These patient feature representations were used to associate clinical information within each dataset to evaluate the overall methodology.

Patient groups were identified by k-Means clustering on patient-level feature representations. The optimal number of clusters was determined using the elbow method with inertia. Inertia is the sum of all element distances to the cluster center in one cluster. It decreases when the number of clusters is increased. The optimal number of clusters is the point where the inertia stops decreasing drastically. Then, we calculated the univariate Kaplan-Meier estimator and performed a log-rank test for each patient group. Moreover, in the multivariate Cox Proportional Hazards Model, the group with the best survival estimation in the Kaplan-Meier plot is used as baseline for verification. We used lifelines^48^ library for the survival analyses.

### Creating attention maps of cores

Attention maps were generated by inputting an image into the ViT model, and extracting [CLS] attention heads from different layers independently to capture the focus of the model on the image. To create core-level attention maps, we used the local models. Firstly, we cropped the input core image into overlapping small patches of size 256 × 256 pixels, with an overlap of 128 pixels. Then, we input those overlapped patches to the local ViT model, and extracted the attention maps for each patch. The attention maps were stitched back together based on the original positions of the respective input patches. Then, the values of the overlapping areas were averaged. Lastly, we masked out the area outside of the true core with a loose mask created by dilating the DAPI channel. For visualizations in Fig. 5, we show only values above the upper quartile to highlight the most relevant regions inside the images.

### Clustering metrics

We used multiple metrics on the feature representation space. All metrics are calculated with cosine distance.

#### Normalized Mean Pairwise Distance (nMPD)

nMPD is the average distance (cosine in our case) between all pairs of points in the set divided by the largest distance. In the context of high dimensional datapoints, nMPD is close to 0 when the datapoints are similar to each other. On the other hand, nMPD is closer to 1 when the datapoints are sufficiently far away from all other datapoints.

#### Silhouette score

To evaluate separation of local features from different regions, we used average Silhouette score. Average Silhouette score of 0 or lower indicates poor clustering with complete overlap of clusters, and a score of 1 indicates perfect separation between clusters with regards to the label.

#### Classification scores

We calculated precision, recall, specificity, and F1 scores using kNN classification of tumor regions, while considering samples from the tumor-center region as positive. To evaluate the feature representations thoroughly, the classification scores are calculated with different numbers of neighbors for the kNN. Ideally, classification scores should be high with a smaller number of neighbors and gradually decrease as the number of neighbors increases. This should indicate that immediate neighboring points are similar to each other, while a larger neighborhood does not need to exhibit the same level of similarity.

### Visualizations

Core visualizations show values between the median and the 99th percentile for each channel. Values below a lower threshold are set to zero to remove the background, while values above an upper threshold are set to the upper threshold value. However, inputs to the models are always logarithmic transformations of the raw images. Uniform Manifold Approximation and Projection (UMAP) for dimension reduction visualizations are created by first reducing the dimensionality to 50 using principal component analysis (PCA) followed by computing the two-dimensional UMAP for visualization.

## Data availability

Source code is available on GitHub repository: https://github.com/bioimage-profiling/SSL-Multiplexed-Imaging.

The trained MAE encoders of the local models are available in the Zenodo repository: https://zenodo.org/records/15011375

## Acknowledgement

- We acknowledge the Research Council of Finland (grants 340273, 346604, 359907, 363154 for L.P.; 363152 for T.P.; 322675 for T.M.), the Cancer Foundation Finland (220040 for L.P.; decision date 30/11/2022 for T.P.; 304667, 191118 for T.M.), Jane and Aatos Erkko Foundation (290520 for T.M.), Hospital District of Helsinki and Uusimaa (TYH2018214, TYH2018222, TYH2019235, TYH2019249 for T.M.), and Sigrid Juselius Foundation (230149, 240158 for T.P.; 8191 for T.M.)
- We want to acknowledge the participants and investigators of iCAN study. The iCAN project is funded by the Research Council of Finland (grant number 320185), University of Helsinki, Helsinki University Hospital and industry partner Boehringer Ingelheim.
- We acknowledge CSC–IT Center for Science, Finland for awarding this project access to the LUMI supercomputer, owned by the EuroHPC Joint Undertaking, hosted by CSC (Finland) and the LUMI consortium.
- Microscopic imaging of the samples was performed at the FIMM Digital Microscopy and Molecular Pathology Unit, supported by HiLIFE and Biocenter Finland.
- Johan Botling and Patrick Micke are acknowledged for lung cancer cohort clinical data.
- We acknowledge Anna Nylund for help with the visual appearance of figures.

## Author information

### Authors and Affiliations

**Institute for Molecular Medicine Finland (FIMM), HiLIFE, University of Helsinki, Helsinki, 00014, Finland**

Gantugs Atarsaikhan, Isabel Mogollon, Katja Välimäki, Teijo Pellinen & Lassi Paavolainen

**iCAN Digital Precision Cancer Medicine Flagship, University of Helsinki, Helsinki, 00014, Finland**

Gantugs Atarsaikhan, Katja Välimäki & Lassi Paavolainen

**Department of Pathology, University of Helsinki and Helsinki University Hospital, Helsinki, 00029, Finland**

Tuomas Mirtti

**Research Program in Systems Oncology, University of Helsinki, Helsinki, 00014, Finland**

Tuomas Mirtti

**Finnish Cancer Institute, Helsinki, Finland**

Tuomas Mirtti

### Consortia

**iCAN**

Names of the iCAN Digital Precision Cancer Medicine Flagship project members are provided in the Supplementary Information.

### Contributions

G.A. and L.P. conceived and designed the study. G.A. performed data preprocessing, implemented the self-supervised learning framework, and performed the data analysis. K.V. stained and imaged tissue samples. I.M. and K.V. contributed to data preprocessing and result interpretation. T.M. provided clinical samples, pathology expertise, and assisted in clinical interpretation. G.A., L.P. and T.P. wrote the manuscript. L.P. and T.P. supervised the project and contributed to the overall study design and interpretation. All authors contributed to manuscript writing, critically reviewed the content, and approved the final version.

