## Supplementary_material for "Self-supervised learning enables unbiased patient characterization from multiplexed cancer tissue microscopy images"

#### Supplementary methods

##### Benchmarking self-supervised learning methods for multiplexed immunofluorescence microscopy

The proposed framework is designed as modular so that various SSL methods can be used as local and global feature learners. We experimented with four SSL strategies as local learners: DINO, MAE, SimCLR, and VICRegL; and with two SSL strategies as global learners: DINO and MAE.

First, we compared local feature representations of the four approaches with the prostate cancer dataset. All of the approaches learn similar features that can differentiate between core regions, such as *tumor-adjacent* and *tumor-center*, although significant overlap exists between the regions (Suppl. Fig. S1). All methods result in relatively high F1-score (0.63–0.74) but lower specificity score (Suppl. Fig. S2A) when tested with class-balanced k-nearest neighbor (kNN) classification while setting the *tumor-center* as positive (Suppl. Fig. S2B). We hypothesize that the models have high confidence in classifying *tumor-center* patches correctly since patches from those spots contain mostly cancer cells. However, the models are not as confident in classifying *tumor-adjacent* patches correctly due to similar cancer cells included in both regions. Moreover, F1-scores were larger with lower number of neighbors in kNN classifications, indicating that feature representations of patches from the same region are closer together in the feature dimension space. DINO and MAE result in the highest F1-scores, whereas SimCLR, arguably, results in the lowest F1-score and specificity (Suppl. Fig. S2A).

Next, we compared global feature representations of prostate cancer dataset using DINO and MAE as the global SSL methods. The feature representations show more distinct grouping when the global method is DINO rather than MAE (Suppl. Fig. S1). All eight combinations are still able to differentiate between core regions. Using MAE as a global method, F1-scores are systematically similar with the increasing numbers of neighbors, indicating there are no clear differences between cores that are captured (Suppl. Fig. S2C). Contrary, when DINO is used as the global method, the kNN F1-scores are decreasing with increasing number of neighbors, indicating improved grouping of similar cores. DINO-DINO combination (local: DINO and global: DINO) is able to distinguish between tumor-centers, however, it struggles to correctly classify tumor-adjacent spots with low specificity (Suppl. Fig. S2C). MAE-DINO and VICRegL-DINO combinations are better at classifying both tumor-center and tumor-adjacent spots.

We then looked into batch effects in the feature representations (Suppl. Fig. S2B, D). The prostate cancer dataset includes nine TMA slides with approximately hundred spots each. We defined each TMA slide as a separate batch and calculated kNN scores in two ways: leave-one-slide-out strategy, by ignoring patches or spots from the same slide when testing kNN, and without introducing any restrictions. We studied the difference between kNN scores of leave-one-slide-out and no-restriction classifications. In local, SimCLR local feature representations show the smallest batch effects, although it also had the lowest scores on classification tasks (Suppl. Fig. S2B). Other methods suffer similarly from batch effects, however, having higher scores on classification tasks. In global, kNN score differences are small compared to local with VICRegL-DINO and MAE-DINO combinations having the smallest difference (Suppl. Fig. S2D). In general, the differences between restricted and non-restricted classifications are negligible, indicating that TMA slide level batch effect is minimal.

Lastly, we determined the local model training epochs based on the UMAP visualizations of feature representations (Suppl. Fig. S3). Stopping the training too early (epoch=10) led to incomplete learning of features that appeared as a structure without any visual clusters. On the other hand, visualizations from later epochs (epoch=200) showed that patches from the same cores were grouping together strongly which appeared as an extreme batch-effect. This indicates that the model learned features only specific to each core rather than meaningful representation of the whole dataset. We selected models after an epoch when the UMAP visualization started to overfit to individual cores.

Overall, the local feature representations generated by the DINO method were the most expressive, followed by those from the MAE method. However, at global, methods trained using MAE local feature representations produced more meaningful groupings. The global models trained on VICRegL local feature representations as inputs were also effective, although VICRegL local feature representations were less expressive. Thus, for further analysis, we used feature representations of the MAE-DINO combination.

#### Supplementary Figure S1

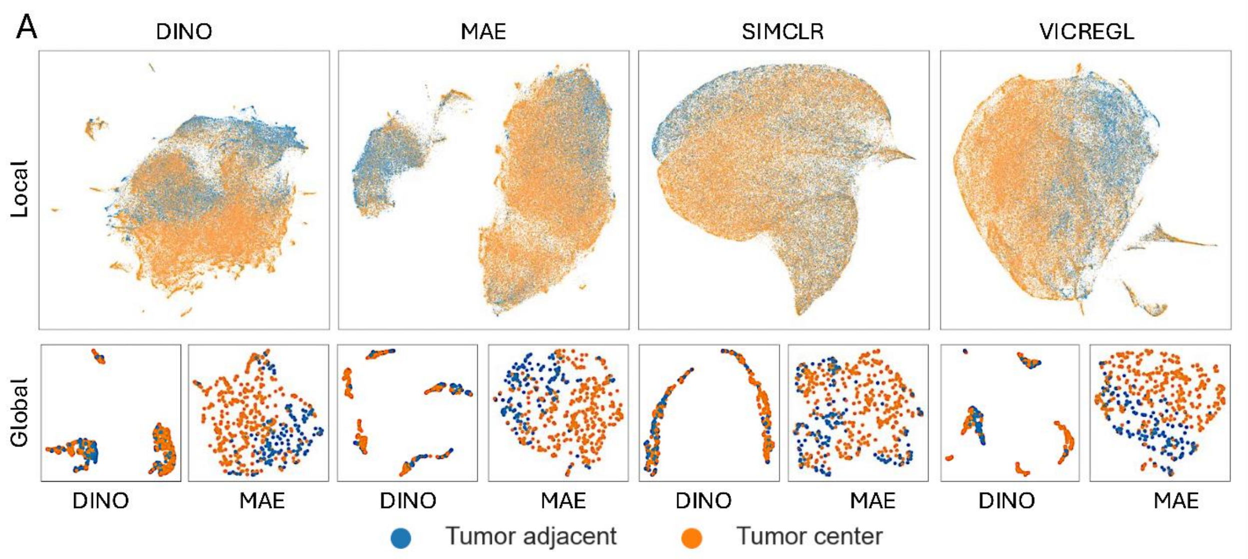


**Supplementary Figure S1. Comparison of self-supervised learning approaches on prostate cancer cohort.** Top row: UMAPs show local feature representations of DINO, MAE, SimCLR, and VICRegL SSL methods when trained on the prostate cancer dataset. Each dot represents a patch from tumor-center (orange) or from tumor-adjacent (blue) core. Bottom row: UMAPs show global feature representations of combinations using DINO or MAE as a global method on the corresponding local feature representations.

### Supplementary Figure S2

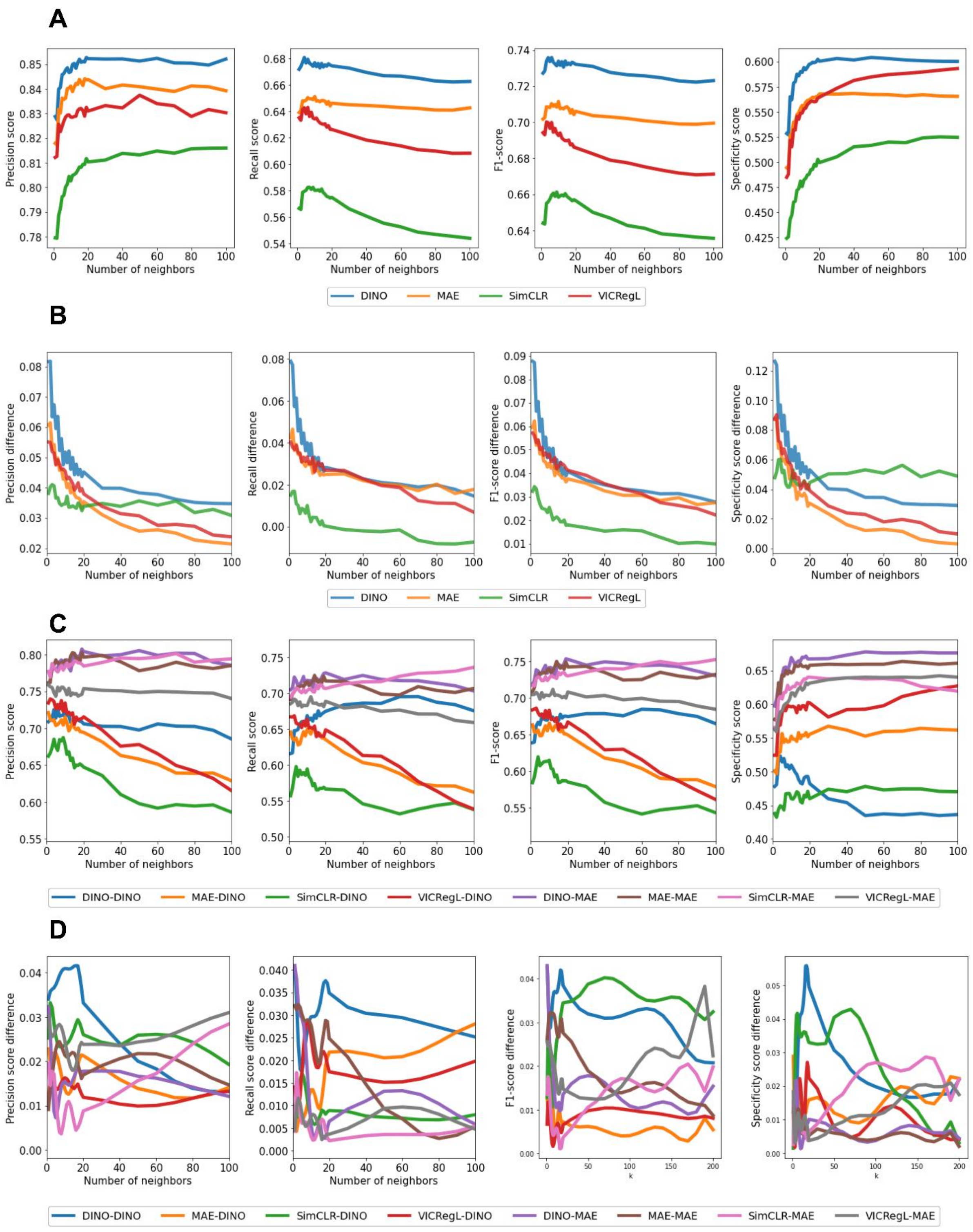


**Supplementary Figure S2. Tumor region classification performances using different methods.** The graphs show the classification scores for the tumor region classification task using k-nearest neighbor (kNN) as *k* is gradually increased with the prostate cancer cohort. Tumor-center is considered as the positive category. **A)** Scores from local feature representations. We used a *leave-one-patient-out* strategy, meaning that we excluded patches from the same patient from evaluation. **B)** Differences between kNN scores between *leave-one-slide-out* classification and no-restriction classification using local feature representations. Scores are closer to zero meaning that slide-level batch effect is negligible. **C)** Classification scores of *leave-one-patient-out* kNN classification using global features. **D)** Differences between kNN scores between *leave-one-slide-out* classification and no-restriction classification using DINO as the global method.

#### Supplementary Figure S3

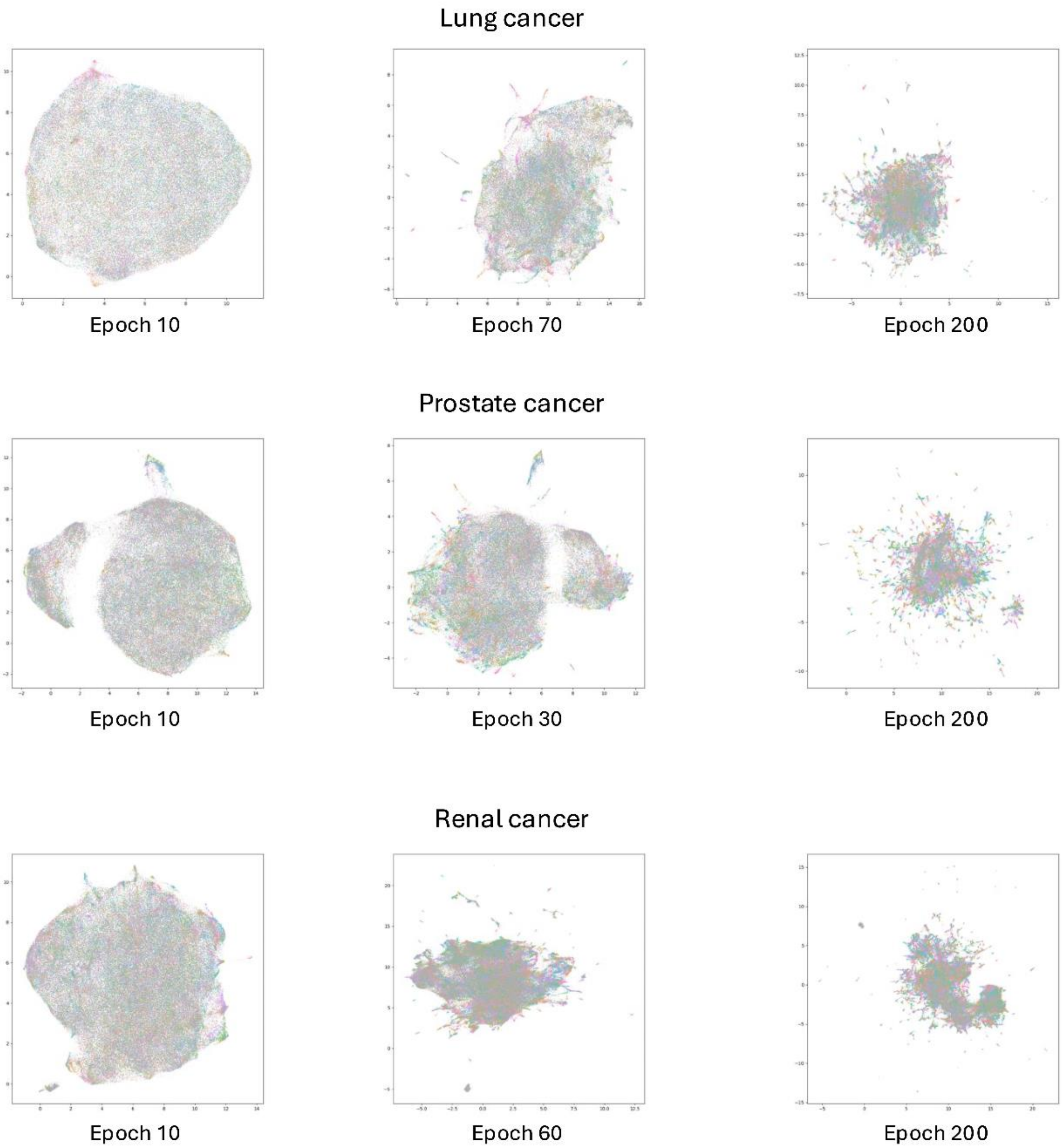


**Supplementary Figure S3. UMAP visualizations of local features trained with the DINO approach.** Each color represents one core. However, since hundreds of cores are presented, some colors are used multiple times. At epoch 10, the DINO model is still underfitted, thus UMAP creates a 'blob'-like structure. When the model is overfitted (epoch 200), the model extremely overfits to the input images and creates one cluster for each core. We stop training when the model just starts to overfit (here epochs 70, 30, and 60).

#### Supplementary Figure S4

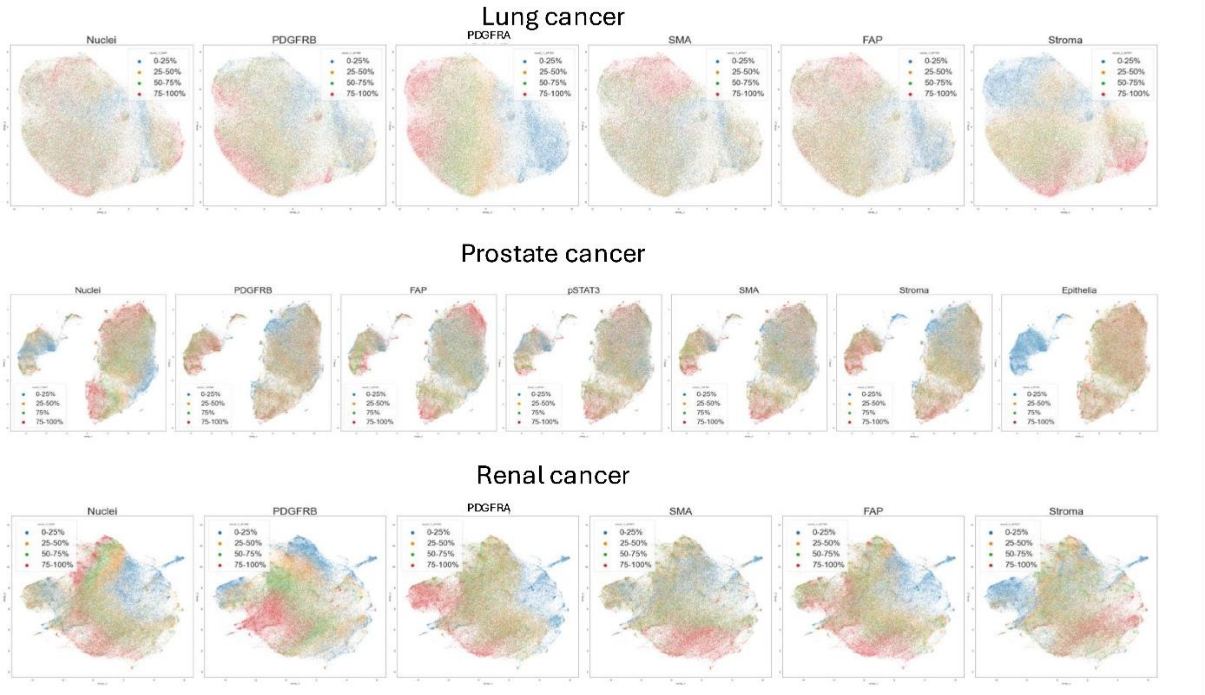


**Supplementary Figure S4. Visualizations of local feature representations on lung, prostate, and renal cancer cohorts.** We calculated mean intensity values of each channel for each patch image. The colors are based on four quartiles. The visualizations show various possible channel combinations captured in the local feature representations.

#### Supplementary Figure S5

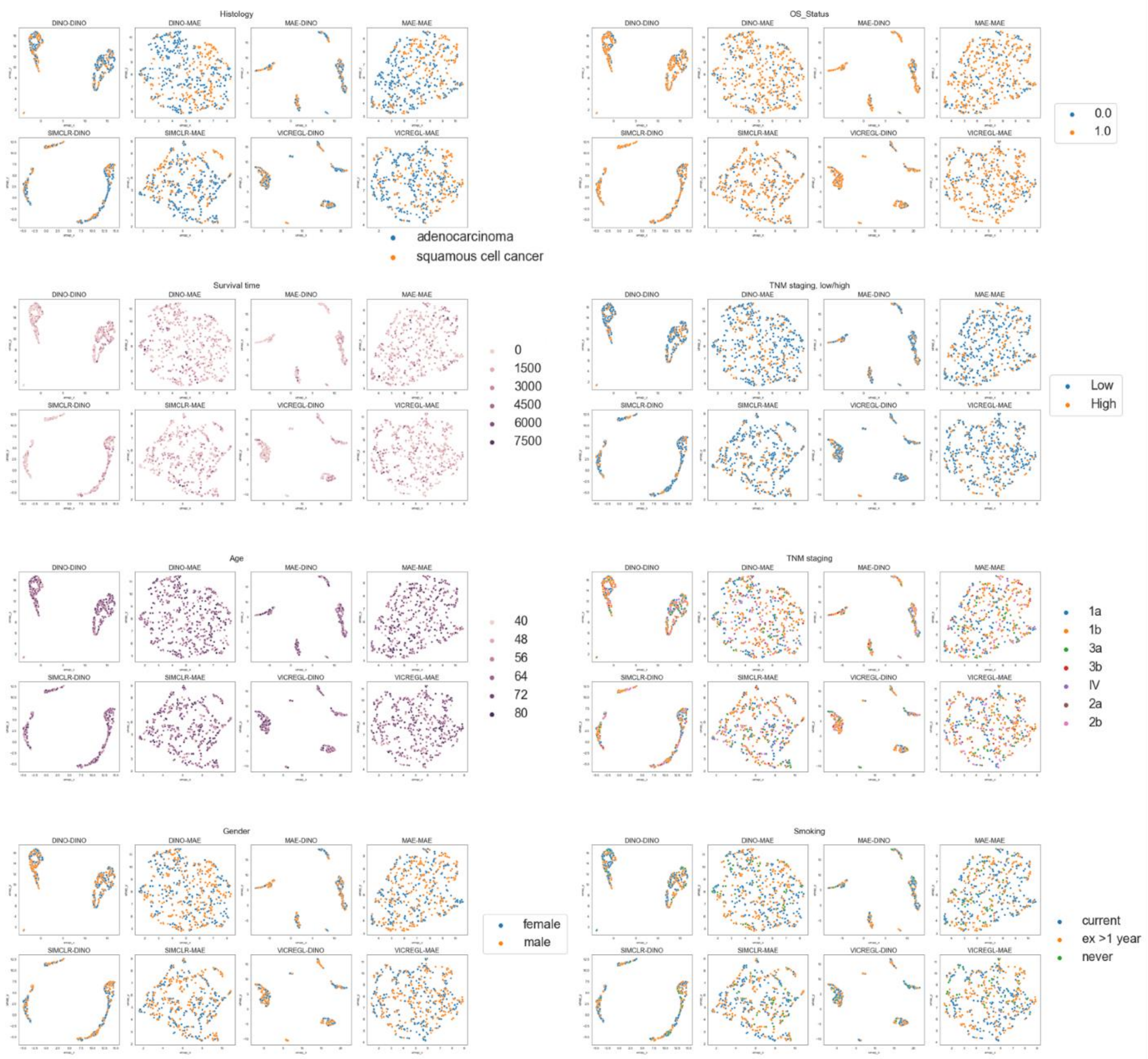


**Supplementary Figure S5. Global feature representations of lung cancer cohort with various clinical information.** All eight combinations of local (DINO, MAE, SimCLR, VICRegL) and global (DINO, MAE) models are plotted.

### Supplementary Figure S6

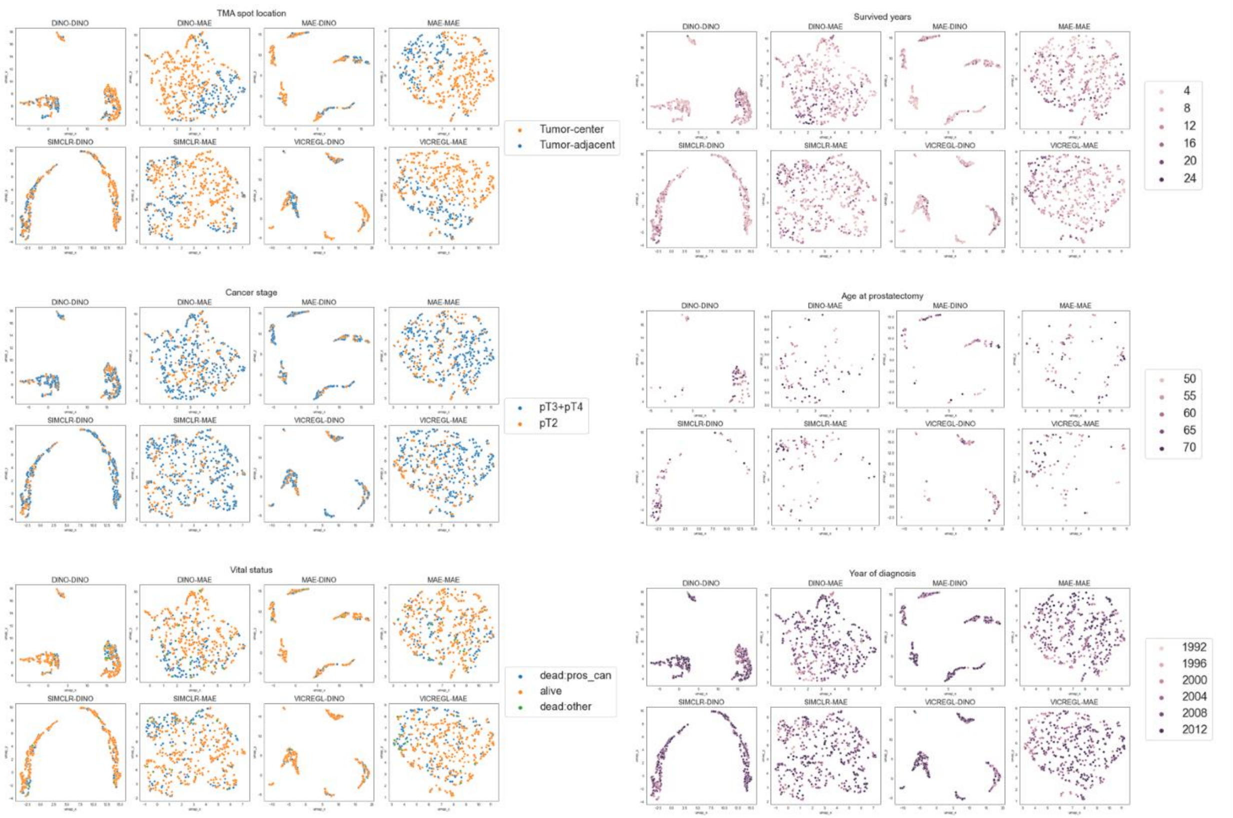


**Supplementary Figure S6. Global feature representations of prostate cancer cohort with various clinical information.** All eight combinations of local (DINO, MAE, SIMCLR, VICRegL) and global (DINO, MAE) models are plotted.

### Supplementary Figure S7

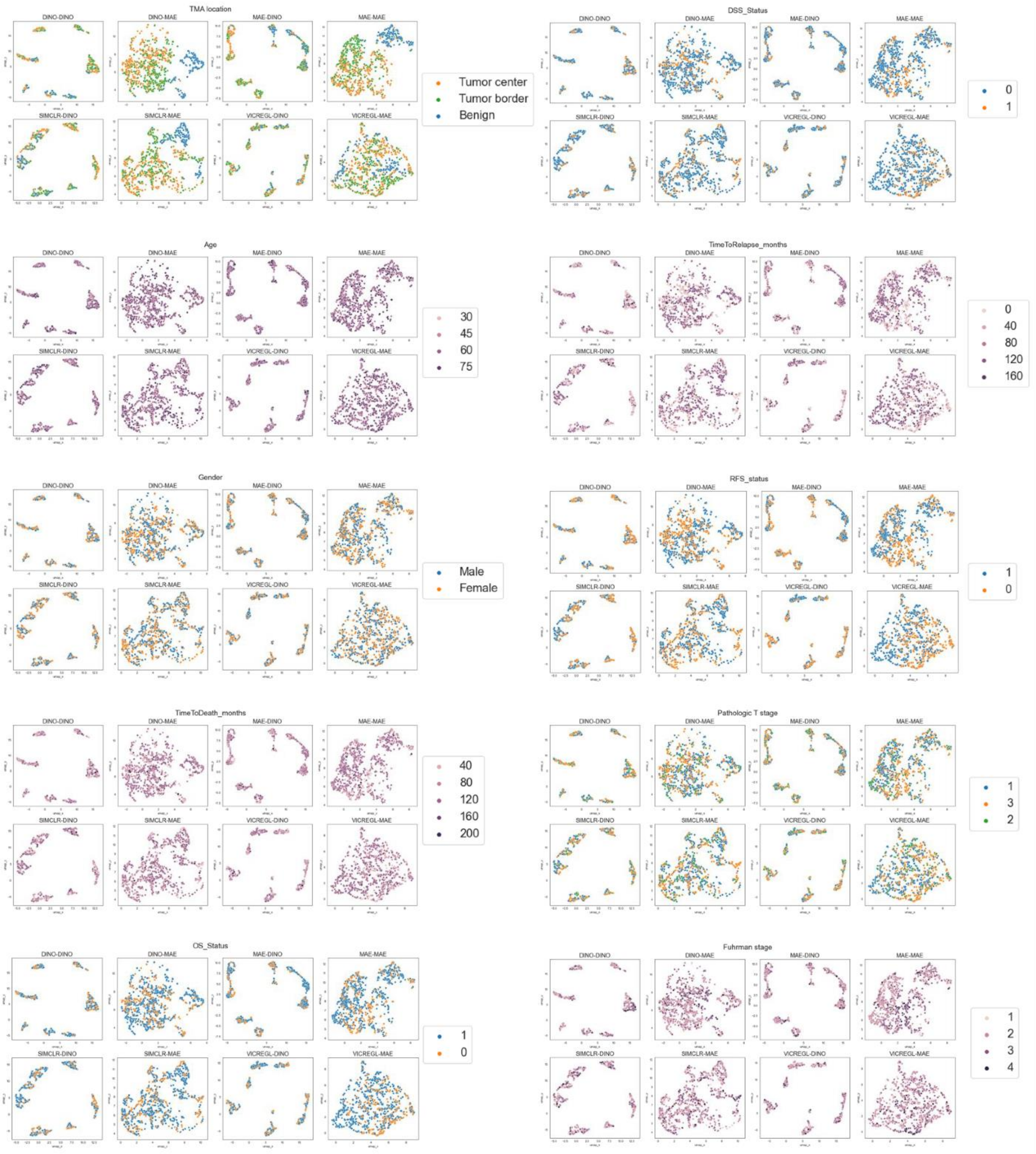


**Supplementary Figure S7. Global feature representations of renal cancer cohort with various clinical information.** All eight combinations of local (DINO, MAE, SIMCLR, VICRegL) and global (DINO, MAE) models are plotted.

### Supplementary Figure S8

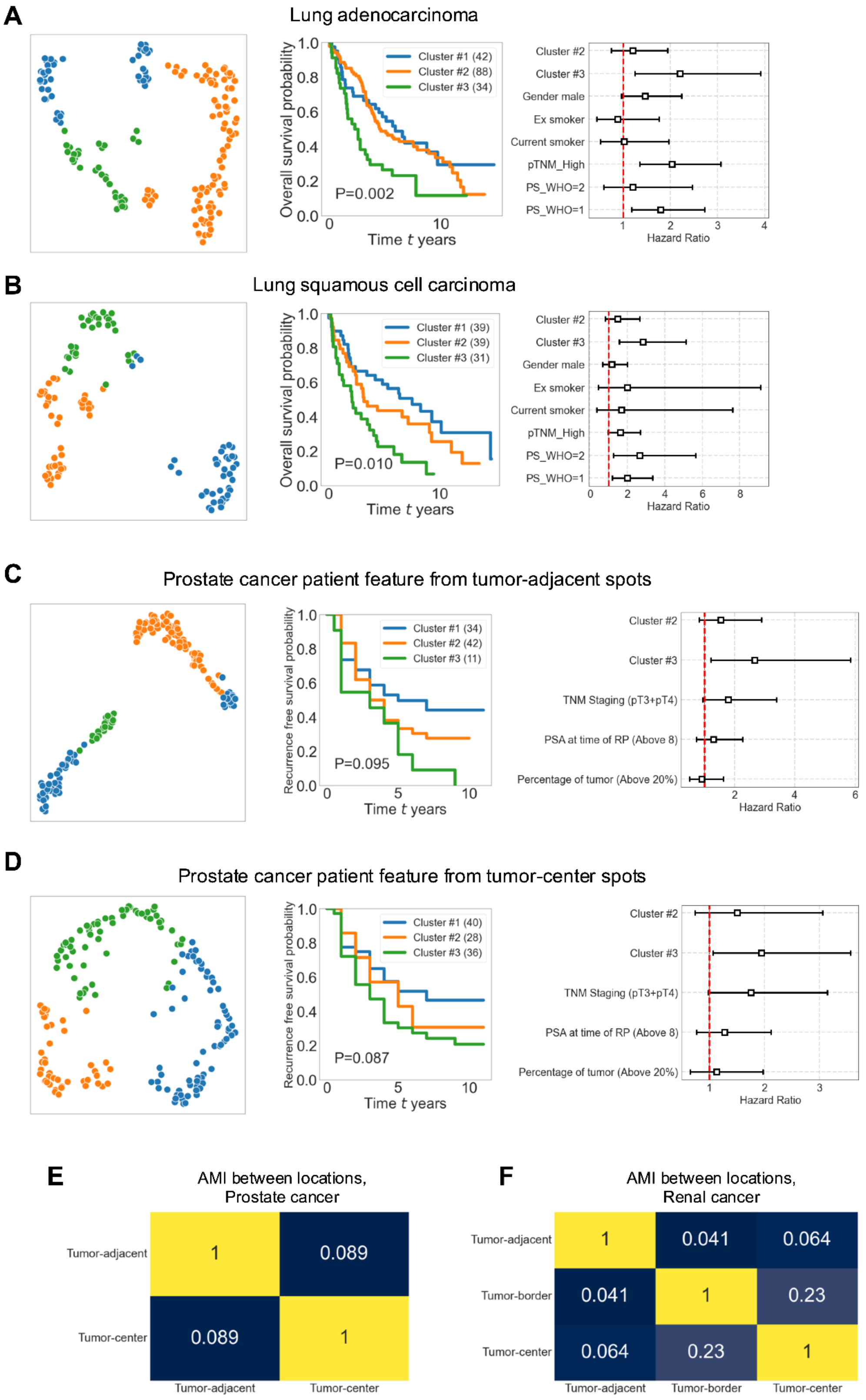

**Supplementary Figure S8. Patient-level feature representations and survival plots for patient groups.** Patient-level feature representations were created individually by averaging all TMA global level features from the same region. We clustered feature representations with k-means clustering. The number of clusters was determined with the elbow method. The color code in graphs (A-D) presents the cluster ID. We studied patient survival or recurrence using Kaplan-Meier plot and calculated the hazard rate using Cox proportional hazard regression. We considered overall survival probability for the lung cancer cohort **(A, B)**, and recurrence-free survival probability following surgical removal of the tumor tissue for the prostate cancer cohort **(C,D)**. Adjusted mutual information (AMI) scores were calculated to study region-wise differences between cluster labels for prostate and renal cancer cohorts**(E, F)**.

### Supplementary Figure S9

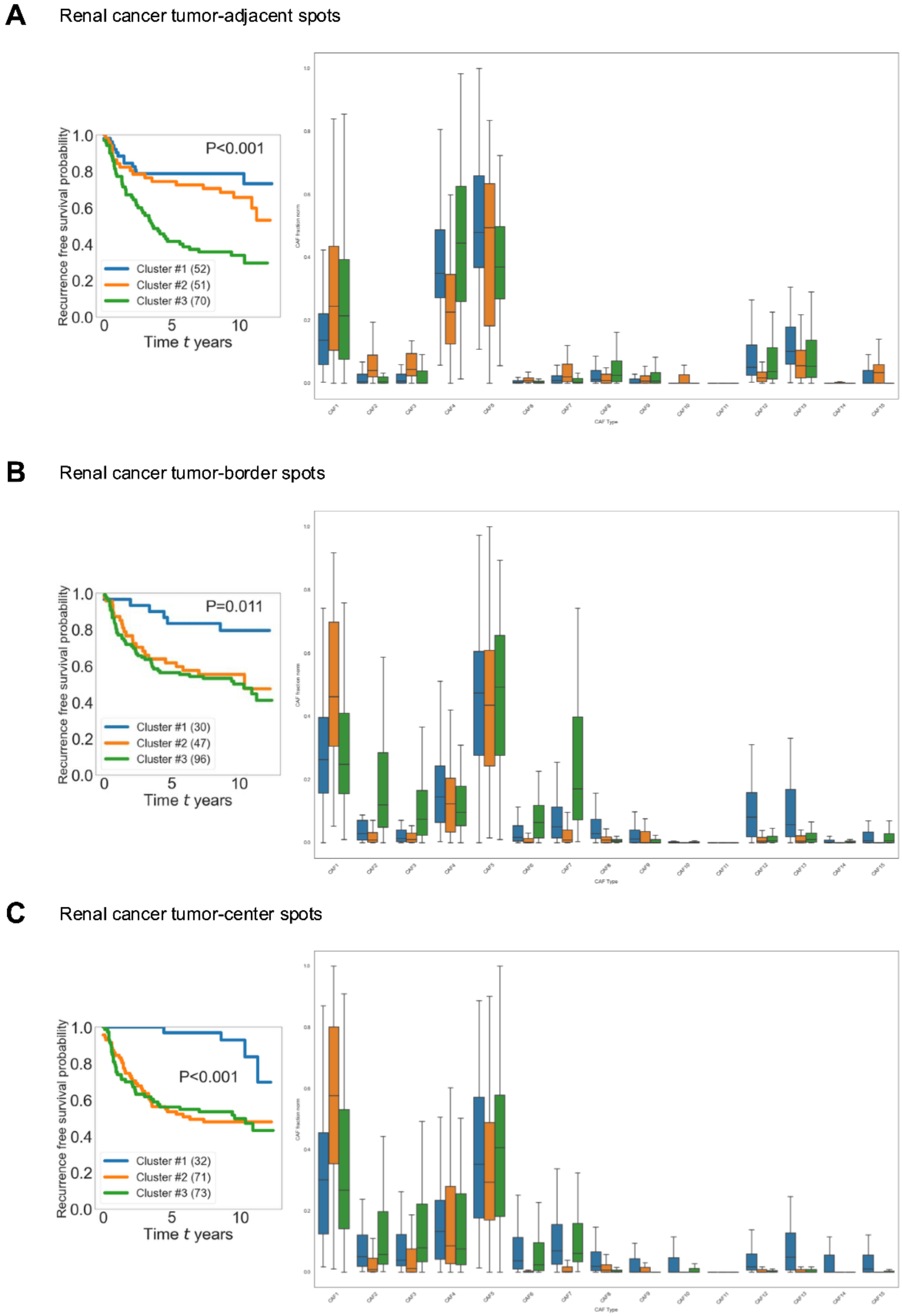


**Supplementary Figure S9. Cancer-associated fibroblast (CAF) associations on renal cancer cohort.** Kaplan-Meier plots for patient clusters determined from patient-level feature representations, with the corresponding CAF ratio (ratio of CAF subtype and all CAFs) in each cluster. CAF fractions are scaled across samples representing different patient clusters for each CAF subtype. Outliers are not shown.

### Supplementary Figure S10

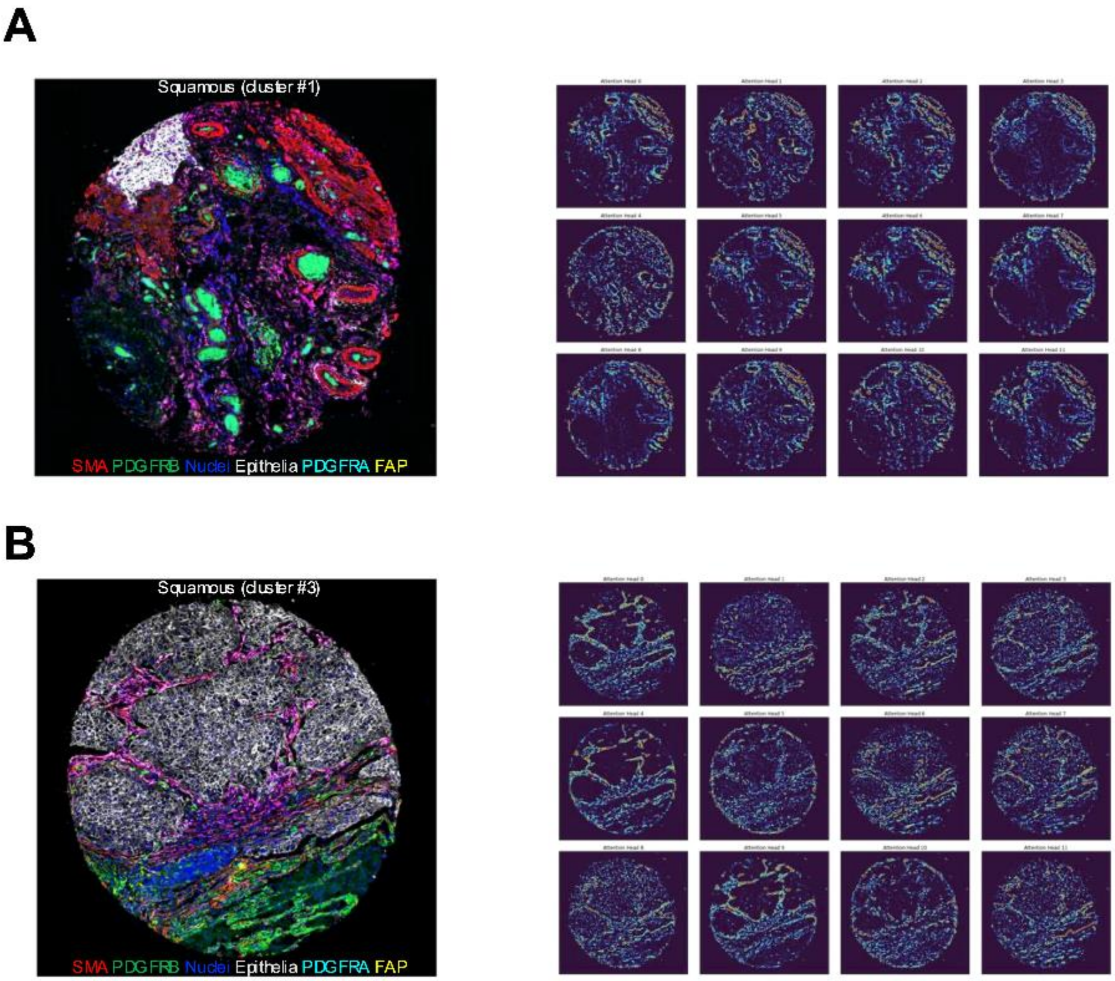


**Supplementary Figure S10. Individual attention maps of lung squamous cell carcinoma cores. A)** An example from good prognosis group (cluster#1), **B)** an example from poor prognosis group (cluster#3). The attention maps are visualized by cutting off values below 80th percentile.

### Supplementary Figure S11

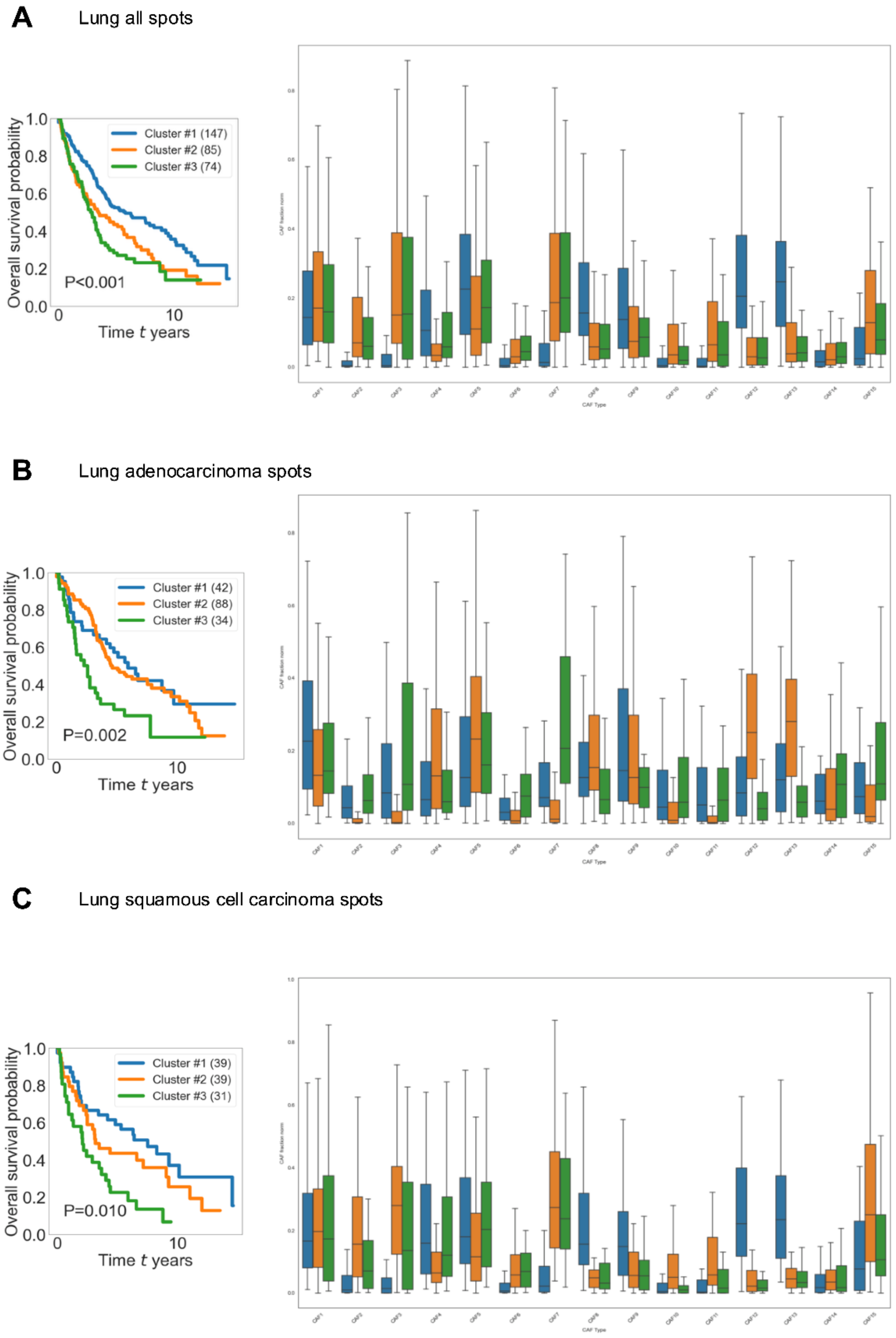


**Supplementary Figure S11. Cancer-associated fibroblast (CAF) associations for lung**

**cancer cohort.** Kaplan-Meier plots for patient clusters determined from patient-level feature representations, with the corresponding CAF ratio (ratio of CAF subtype and all CAFs) in each cluster. CAF fractions are scaled across samples representing different patient clusters for each CAF subtype. Outliers are not shown.

#### Supplementary Table S1

**Supplementary Table S1. CAF subsets.**

| CAF Type | PDGFRB | PDGFRA | SMA | FAP | PanEpiMask |
|---|---|---|---|---|---|
| CAF1 | (-) | (-) | (+) | (-) | (-) |
| CAF2 | (-) | (-) | (-) | (+) | (-) |
| CAF3 | (-) | (-) | (+) | (+) | (-) |
| CAF4 | (+) | (-) | (-) | (-) | (-) |
| CAF5 | (+) | (-) | (+) | (-) | (-) |
| CAF6 | (+) | (-) | (-) | (+) | (-) |
| CAF7 | **(+)** | **(-)** | **(+)** | **(+)** | **(-)** |
| CAF8 | (-) | (+) | (-) | (-) | (-) |
| CAF9 | (-) | (+) | (+) | (-) | (-) |
| CAF10 | (-) | (+) | (-) | (+) | (-) |
| CAF11 | (-) | (+) | (+) | (+) | (-) |
| CAF12 | (+) | (+) | (-) | (-) | (-) |
| CAF13 | **(+)** | **(+)** | **(+)** | **(-)** | **(-)** |
| CAF14 | (+) | (+) | (-) | (+) | (-) |
| CAF15 | (+) | (+) | (+) | (+) | (-) |

### Supplementary Table S2

**Supplementary Table S2. Cluster-wise patient summary of lung cancer cohort.**

| | | Adenocarcinoma Cluster #1 | Adenocarcinoma Cluster #2 | Adenocarcinoma Cluster #3 | Squamous Cluster #1 | Squamous Cluster #2 | Squamous Cluster #3 |
|---|---|---|---|---|---|---|---|
| **patient count** | | **44** | **90** | **34** | **40** | **40** | **31** |
| Metadata | Unique values | | | | | | |
| Gender | female | 25 | 49 | 22 | 11 | 10 | 10 |
| | male | 19 | 41 | 12 | 29 | 30 | 21 |
| Age | 67 or less | 25 | 50 | 17 | 18 | 24 | 20 |
| | Above 67 | 19 | 40 | 17 | 22 | 16 | 11 |
| pTNM 7th | 1a | 15 | 35 | 8 | 8 | 6 | 4 |
| | 1b | 13 | 31 | 15 | 19 | 17 | 17 |
| | 2a | 2 | 4 | 0 | 0 | 1 | 0 |
| | 2b | 8 | 9 | 3 | 7 | 9 | 1 |
| | 3a | 2 | 7 | 4 | 4 | 5 | 5 |
| | 3b | 1 | 1 | 2 | 2 | 1 | 4 |
| | IV | 3 | 3 | 2 | 0 | 1 | 0 |
| pTNM lowVsHigh | 0 | 28 | 66 | 23 | 27 | 23 | 21 |
| | 1 | 16 | 24 | 11 | 13 | 17 | 10 |
| Status | 0 | 17 | 28 | 7 | 16 | 11 | 4 |
| | 1 | 27 | 62 | 27 | 24 | 29 | 27 |
| Smoking | current | 19 | 38 | 19 | 16 | 24 | 12 |
| | ex >1 year | 21 | 36 | 12 | 23 | 16 | 16 |
| | never | 4 | 14 | 3 | 1 | 0 | 3 |
| PS WHO | 0 | 28 | 57 | 19 | 19 | 22 | 11 |
| | 1 | 13 | 26 | 12 | 17 | 14 | 17 |
| | 2 | 2 | 5 | 3 | 3 | 4 | 2 |

| | | | | | | | |
|---|---|---|---|---|---|---|---|
| | 3 | 1 | 1 | 0 | 1 | 0 | 1 |
| | 4 | 0 | 1 | 0 | 0 | 0 | 0 |
| adj treatm | not recorded | 24 | 58 | 19 | 16 | 23 | 20 |
| | no | 14 | 25 | 9 | 20 | 8 | 9 |
| | yes | 6 | 7 | 6 | 4 | 9 | 2 |

#### Supplementary Table S3

**Supplementary Table S3. Cluster-wise patient summary of prostate cancer cohort.**

| | | Tumor-adjacent Cluster #1 | Tumor-adjacent Cluster #2 | Tumor-adjacent Cluster #3 | Tumor-center Cluster #1 | Tumor-center Cluster #2 | Tumor-center Cluster #3 |
|---|---|---|---|---|---|---|---|
| **patient count** | | **57** | **72** | **19** | **68** | **44** | **61** |
| Metadata | Unique values | | | | | | |
| Post op. hormone therapy | No | 54 | 68 | 17 | 65 | 43 | 56 |
| | Yes | 3 | 4 | 2 | 3 | 1 | 5 |
| Previous history of other cancer | No | 55 | 70 | 18 | 64 | 44 | 59 |
| | Unknown | 2 | 1 | 1 | 3 | 0 | 2 |
| | Yes | 0 | 1 | 0 | 1 | 0 | 0 |
| Biopsy Gleason score | 4 | 1 | 1 | 0 | 0 | 1 | 1 |
| | 5 | 1 | 3 | 2 | 1 | 4 | 2 |
| | 6 | 9 | 14 | 0 | 10 | 9 | 11 |
| | 7 | 34 | 38 | 5 | 39 | 20 | 31 |
| | 8 | 5 | 9 | 8 | 8 | 6 | 11 |
| | 9 | 6 | 2 | 3 | 7 | 1 | 3 |
| PSA at time of diagnosis (ng/ml) | 9 or less | 29 | 47 | 10 | 34 | 27 | 37 |
| | Above 9 | 28 | 25 | 9 | 34 | 17 | 24 |
| PSA at time of RP | 9 or less | 23 | 37 | 10 | 25 | 25 | 31 |
| | Above 9 | 34 | 35 | 9 | 43 | 19 | 30 |
| Prostate wet weight (g) | 48 or less | 36 | 36 | 10 | 34 | 24 | 34 |
| | Above 48 | 21 | 36 | 9 | 34 | 20 | 27 |
| TNM 8. Ed. Staging | pT2 | 14 | 25 | 4 | 17 | 22 | 16 |
| | pT3+pT4 | 43 | 47 | 15 | 51 | 22 | 45 |
| Lymph Nodes Removed at RP | No | 9 | 21 | 2 | 16 | 4 | 15 |
| | Removed | 0 | 1 | 0 | 0 | 0 | 1 |

| | | | | | | | |
|---|---|---|---|---|---|---|---|
| | before | | | | | | |
| | Unknown | 22 | 6 | 5 | 30 | 8 | 8 |
| | Yes | 26 | 44 | 12 | 22 | 32 | 37 |
| RP/TURP Gleason score (6, 7, 8, 9, 10) | 7 | 44 | 57 | 11 | 50 | 39 | 43 |
| | 8 | 3 | 12 | 4 | 4 | 5 | 14 |
| | 9 | 9 | 3 | 4 | 13 | 0 | 4 |
| | 10 | 1 | 0 | 0 | 1 | 0 | 0 |
| Percentage of cancer (of whole prostate), number (%) | 20 or less | 30 | 53 | 12 | 36 | 39 | 39 |
| | Above 20 | 27 | 19 | 7 | 32 | 5 | 22 |

#### Supplementary Table S4

**Supplementary Table S4. Cluster-wise patient summary of renal cancer cohort.**

| | | Tumor-adjacent Cluster #1 | Tumor-adjacent Cluster #2 | Tumor-adjacent Cluster #3 | Tumor-border Cluster #1 | Tumor-border Cluster #2 | Tumor-border Cluster #3 | Tumor-center Cluster #1 | Tumor-center Cluster #2 | Tumor-center Cluster #3 |
|---|---|---|---|---|---|---|---|---|---|---|
| **Patient count** | | **52** | **53** | **71** | **30** | **47** | **99** | **32** | **71** | **76** |
| Metadata | Unique values | | | | | | | | | |
| Age | 65 or less | 30 | 29 | 29 | 13 | 21 | 54 | 15 | 35 | 40 |
| | Above 65 | 22 | 24 | 42 | 17 | 26 | 45 | 17 | 36 | 36 |
| Gender | Female | 25 | 26 | 35 | 13 | 27 | 43 | 10 | 34 | 42 |
| | Male | 27 | 27 | 36 | 17 | 20 | 56 | 22 | 37 | 34 |
| OS_Status | 0 | 43 | 36 | 38 | 26 | 31 | 60 | 28 | 48 | 43 |
| | 1 | 9 | 16 | 33 | 4 | 16 | 38 | 4 | 23 | 32 |
| DSS_Status | 0 | 46 | 40 | 43 | 28 | 35 | 66 | 31 | 52 | 48 |
| | 1 | 6 | 12 | 27 | 2 | 11 | 32 | 1 | 18 | 27 |
| RelapseLocalization | 0 | 42 | 36 | 26 | 26 | 27 | 51 | 31 | 36 | 39 |
| | lung | 6 | 8 | 36 | 1 | 15 | 34 | 1 | 22 | 28 |

| | | | | | | | | | | |
|---|---|---|---|---|---|---|---|---|---|---|
| | liver | 1 | 4 | 0 | 0 | 1 | 4 | 0 | 2 | 3 |
| | brain | 2 | 0 | 1 | 0 | 1 | 2 | 0 | 2 | 1 |
| | kidney | 1 | 2 | 6 | 2 | 2 | 5 | 0 | 6 | 3 |
| RFS_status | 0 | 40 | 32 | 24 | 24 | 25 | 47 | 28 | 34 | 36 |
| | 1 | 12 | 20 | 47 | 6 | 22 | 51 | 4 | 37 | 39 |
| pT_simple | 1 | 31 | 26 | 24 | 19 | 19 | 43 | 23 | 25 | 34 |
| | 2 | 6 | 4 | 13 | 4 | 6 | 13 | 2 | 11 | 11 |
| | 3 | 15 | 23 | 34 | 7 | 22 | 43 | 7 | 35 | 31 |
| Fuhrman stage | 1 | 5 | 3 | 3 | 3 | 2 | 5 | 4 | 6 | 1 |
| | 2 | 32 | 30 | 36 | 19 | 30 | 51 | 23 | 39 | 39 |
| | 3 | 13 | 17 | 27 | 8 | 13 | 36 | 5 | 24 | 28 |
| | 4 | 2 | 3 | 5 | 0 | 2 | 7 | 0 | 2 | 8 |

#### Supplementary Table S5

**Supplementary Table S5. Cluster-wise patient summary of lung cancer cohort (including both histologies).**

| | | Cluster #1 | Cluster #2 | Cluster #3 |
|---|---|---|---|---|
| **Patient count** | | 152 | 88 | 75 |
| Metadata | Unique values | | | |
| Gender | female | 73 | 42 | 27 |
| | male | 79 | 46 | 48 |
| Age | 67 or less | 83 | 49 | 43 |
| | Above 67 | 69 | 39 | 32 |
| pTNM_7th | 1a | 46 | 24 | 12 |
| | 1b | 62 | 35 | 37 |
| | 2a | 5 | 1 | 1 |
| | 2b | 18 | 12 | 10 |
| | 3a | 13 | 10 | 7 |
| | 3b | 4 | 1 | 6 |
| | IV | 3 | 5 | 2 |
| pTNM_lowVsHigh | 0 | 108 | 59 | 49 |
| | 1 | 44 | 29 | 26 |
| Status | 0 | 55 | 22 | 17 |
| | 1 | 96 | 66 | 58 |
| Smoking | current | 64 | 46 | 36 |
| | ex >1 year | 70 | 37 | 32 |
| | never | 16 | 5 | 6 |
| PS_WHO | 0 | 83 | 52 | 33 |
| | 1 | 56 | 27 | 35 |
| | 2 | 9 | 9 | 5 |
| | 3 | 3 | 0 | 1 |
| | 4 | 0 | 0 | 1 |
| adj_treatm | nan | 86 | 46 | 49 |
| | no | 51 | 27 | 16 |

| | | | | |
|---|---|---|---|---|
| | yes | 15 | 15 | 10 |

#### Supplementary Table S6

**Supplementary Table S6. Cluster-wise patient summary of prostate cancer cohort (patient-wise concatenated features of both regions).**

| | | Cluster #1 | Cluster #2 | Cluster #3 |
|---|---|---|---|---|
| **Patient count** | | 35 | 44 | 63 |
| Metadata | Unique value | | | |
| Post op. hormone therapy | No | 34 | 41 | 58 |
| | Yes | 1 | 3 | 5 |
| Previous history of other cancer | No | 33 | 42 | 62 |
| | Unknown | 1 | 2 | 1 |
| | Yes | 1 | 0 | 0 |
| Biopsy Gleason score | 4 | 0 | 1 | 1 |
| | 5 | 1 | 0 | 5 |
| | 6 | 3 | 6 | 13 |
| | 7 | 23 | 27 | 23 |
| | 8 | 4 | 3 | 15 |
| | 9 | 2 | 6 | 3 |
| PSA at time of diagnosis (ng/ml) | 11 or less | 15 | 18 | 39 |
| | Above 11 | 20 | 26 | 24 |
| PSA at time of RP | 9 or less | 12 | 14 | 38 |
| | Above 9 | 23 | 30 | 25 |
| Prostate wet weight (g) | 49 or less | 18 | 21 | 32 |
| | Above 49 | 17 | 23 | 31 |
| TNM 8. Ed. Staging | pT2 | 9 | 8 | 21 |
| | pT3+pT4 | 26 | 36 | 42 |
| Lymph Nodes Removed at RP | No | 15 | 6 | 11 |

| | | | | |
|---|---|---|---|---|
| | Removed before | 0 | 0 | 1 |
| | Unknown | 4 | 18 | 6 |
| | Yes | 16 | 20 | 45 |
| RP/TURP Gleason score | 7 | 28 | 32 | 46 |
| | 8 | 4 | 2 | 13 |
| | 9 | 3 | 9 | 4 |
| | 10 | 0 | 1 | 0 |
| Percentage of cancer (of whole prostate), number (%) | 20 or less | 15 | 20 | 38 |
| | Above 20 | 20 | 24 | 25 |

#### Supplementary Table S7

**Supplementary Table S7. Cluster-wise patient summary of renal cancer cohort (patient-wise concatenated features of all regions).**

| | | Cluster #1 | Cluster #2 | Cluster #3 |
|---|---|---|---|---|
| **Patient count** | | 22 | 75 | 76 |
| Metadata | Unique values | | | |
| Age | 65 or less | 13 | 41 | 32 |
| | Above 65 | 9 | 34 | 44 |
| Gender_woman1 | Female | 11 | 36 | 36 |
| | Male | 11 | 39 | 40 |
| OS_Status | 0 | 20 | 55 | 40 |
| | 1 | 2 | 19 | 36 |
| DSS_Status | 0 | 21 | 59 | 47 |
| | 1 | 1 | 15 | 28 |
| RelapseLocalization | 0 | 19 | 55 | 28 |
| | lung | 2 | 10 | 37 |
| | liver | 0 | 3 | 2 |
| | brain | 0 | 1 | 2 |
| | kidney | 1 | 3 | 5 |
| RFS_status2 | 0 | 18 | 50 | 26 |
| | 1 | 4 | 24 | 50 |
| pT_simple | 1 | 14 | 42 | 24 |
| | 2 | 4 | 6 | 12 |
| | 3 | 4 | 27 | 40 |
| Fuhrman stage | 1 | 2 | 5 | 3 |
| | 2 | 14 | 42 | 41 |
| | 3 | 6 | 23 | 28 |
| | 4 | 0 | 5 | 4 |
